# Resource-State Modulation of Prefrontal Value Processing during Earning–Spending Decision Making

**DOI:** 10.64898/2026.11.22.768454

**Authors:** Hualei Wang, Hangyu Si, Lusha Zhu, Tianming Yang

## Abstract

Money mediates earning and spending, and accumulated monetary holdings constitute both a basic form of wealth and an external resource state that shapes economic decisions. Unlike internal physiological states such as hunger or satiety, external resources persist outside the body, can be acquired or depleted through choice, and preserve access to future consumption. Although money and wealth are human constructs, balancing resource acquisition and consumption may rely on evolutionarily conserved neural computations. Here we investigated these computations using an earning–spending task, in which rhesus monkeys chose between earning virtual tokens and spending accumulated tokens for water, with holdings carrying across trials. As holdings varied, monkeys shifted between earning and spending choices. A state-dependent utility model explained this shift through opposing changes in the relative values of earning and spending. We recorded single-unit activities from the anterior cingulate cortex (ACC), orbitofrontal cortex (OFC) and dorsolateral prefrontal cortex (DLPFC). Neuronal populations across all three regions represented current holdings, while ACC and OFC updated these representations after earning and spending outcomes. Crucially, resource state modulated value coding and reshaped the geometry of earning- and spending-value representations across all three regions. OFC exhibited a broader spectrum of signals spanning option value, chosen value, and good-based choice, whereas DLPFC representations were weighted more strongly toward action-based choice. Together, these findings identify distributed prefrontal mechanisms that represent, update, and use an external resource state to guide valuation and choice, extending neural accounts of state-dependent valuation beyond internal physiological conditions and suggesting an evolutionarily conserved foundation for balancing resource acquisition and consumption.

## 1 Introduction

Money is a key intermediary in our everyday economic behavior. People work to acquire money, retain it as wealth, and spend it to obtain goods and experiences. Accumulated monetary holdings therefore constitute an external resource state that shapes these behaviors. Earning an additional unit may be especially valuable when holdings are low but less valuable when they are abundant [1], whereas spending may become more attractive as resources accumulate. Economic decisions therefore depend not only on the opportunities currently available, but also on the resource state in which those opportunities are evaluated [2–4].

This dependence creates a recurring computational problem. An individual must represent current holdings, evaluate opportunities to acquire and consume resources in light of those holdings, and select an option that changes the resource state for subsequent decisions. Resource state thus influences valuation and choice, while earning and spending update the resource state itself [5, 6].

This computational problem resembles the state-dependent valuation processes commonly studied in relation to internal physiological states such as hunger, thirst, and satiety [7–9]. In both cases, current state must be integrated with available opportunities to determine their values [10]. External resources, however, differ fundamentally from internal states. They persist outside the body, can be acquired or depleted through choice, and preserve access to future consumption. Although the neural effects of internal motivational states have been studied extensively [7, 9, 11, 12], comparatively little is known about how the brain represents, updates, and uses accumulated external resources. Here, we ask whether monetary holding, as an external resource state, modulates valuation and choice through related neural principles.

At the neural level, signals related to offer value, chosen value, and categorical choice have been identified across prefrontal cortex, including the orbitofrontal cortex (OFC) [13–15], the dorsolateral prefrontal cortex (DLPFC) [16, 17], and the anterior cingulate cortex (ACC) [18, 19]. Token-based studies have also shown that accumulated holdings influence preferences, including decisions involving gains, losses, and risk, and that neural activity can represent token holdings and other wealth-like reference states [20–23]. These findings identify component signals relevant to resource-dependent choice, but do not explain how they operate together. In particular, it remains unknown how current holdings modulate the values of earning and spending opportunities, how resource-state representations are updated after acquisition and consumption, and how the resulting decision is expressed in good-based and action-based coordinates.

Here we developed a new earning–spending task in which rhesus monkeys chose between acquiring virtual tokens and spending accumulated tokens for water. Tokens carried over across trials, and both earning and spending could continue or stop at intermediate levels of holdings, creating a continuously changing resource state without a fixed redemption threshold. As holdings increased, monkeys shifted systematically from earning toward spending, consistent with opposing state-dependent changes in the relative values of the two options. We combined computational modeling with single-neuron recordings from ACC, OFC, and DLPFC to ask how current resource state alters the relative values of earning and spending, how prefrontal populations represent and update this state, and how state-dependent values are transformed into choices and actions. Together, these findings show how distributed prefrontal representations link accumulated resources to valuation and earning–spending choice.

## 2 Results

### 2.1 Token holdings shift earning–spending preferences

Three rhesus macaques were trained on an earning–spending task (Fig. 1a) in which they chose between performing a fixation task for virtual tokens and spending accumulated tokens for water rewards. Each trial paired one of three earning options, distinguished by earning rate (*V*_Earn_, tokens/s), with one of four spending options, distinguished by water amount in exchange for a token (*V*_Spend_, a.u.) (Fig. 1b). Tokens carried over across trials until spent, so accumulated token holdings constituted a continuous external resource state.

**Fig. 1.**
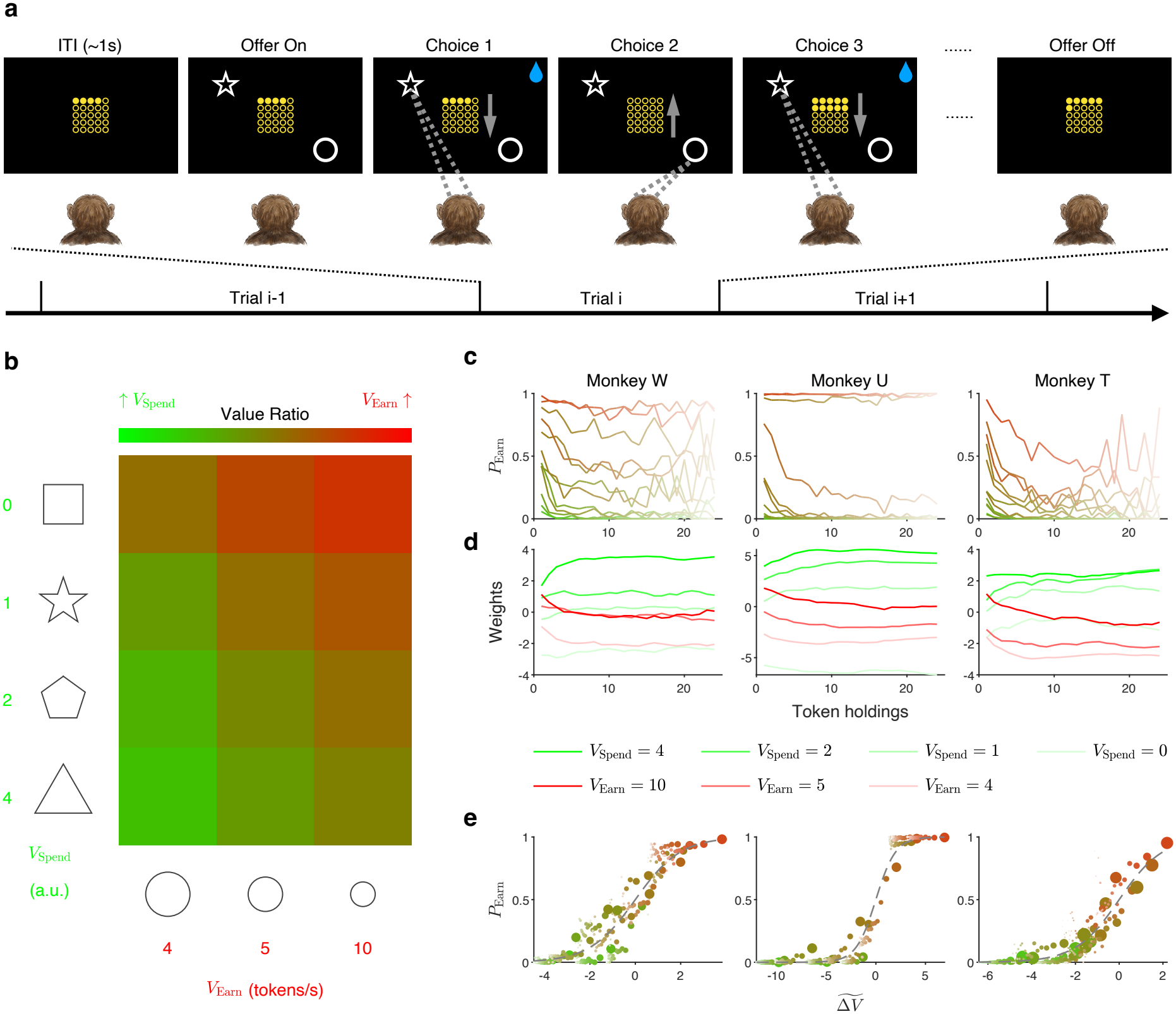
Token holdings shift earning–spending decisions. **(a) Task paradigm.** The central array of yellow circles represents the monkey’s accumulated token holdings (“wallet”), which remained visible during the inter-trial interval (ITI) and operationalized current resource state. Each trial presented one earning and one spending option. Monkeys selected an option by directing their gaze to it and maintaining fixation. During earning, tokens were added to the wallet; during spending, tokens were deducted and water was delivered with each deduction. Monkeys could switch between options by shifting their gaze, and token holdings carried over across trials. **(b) Task conditions**. Twelve (3 × 4) offer conditions were generated by crossing three earning rates (*V*_Earn_ = 4, 5 or 10 tokens/s) with four spending reward magnitudes (*V*_Spend_, a.u.). The spending-reward labels 0, 1, 2 and 4 are schematic (see Methods, Experimental setup and behavioral task). Each earning outcome added one token, whereas each spending outcome cost one token and delivered the corresponding amount of water. The green-to-red gradient schematically orders conditions from those expected to favor spending to those expected to favor earning; it is illustrative and does not represent a quantitatively computed value ratio. **(c) Choice patterns**. Columns show monkeys W, U and T. Earning probability (*P*_Earn_) is plotted against token holdings for each of the 12 offer conditions. Line colors follow the illustrative offer-condition gradient in b, with lighter shades indicating larger token holdings. Only the first choice on each trial was included, provided that it was successfully executed, at holdings of 1–24 tokens. **(d) Subjective weights**. Non-parametric weights for the three earning options (red) and four spending options (green) are plotted against token holdings for each monkey. Color shade indicates earning rate or spending reward magnitude, as shown in the key. **(e) Psychometric curves**. Earning probability is plotted against the normalized value difference Δ*V* = (Δ*V* − *b*)*/γ*, where Δ*V* = *V*_*E*_ − *V*_*S*_ is the difference between the state-dependent earning and spending values, *b* is the fitted choice bias and *γ* is a positive temperature parameter. Smaller *γ* corresponds to greater choice sensitivity. Each point represents a combination of offer condition and token holdings with at least five observations; point size is proportional to the number of observations, and color follows the offer-condition and token-holding conventions in c. Dashed curves show the model-predicted choice probabilities.

The initial choice on each trial revealed two consistent behavioral patterns (Fig. 1c). First, across token holdings, monkeys were more likely to earn when the earning rate was higher and the spending option had less value. This pattern indicates that all three monkeys learned the offer contingencies. Second, within individual offer conditions, the tendency to choose to earn generally declined as tokens accumulated. Thus, choices reflected not only the current offers but also the animal’s current token holdings.

This state-dependent shift in choice is consistent with diminishing marginal utility: accumulating tokens reduces the marginal benefit of earning an additional token and the opportunity cost of spending one. To assess the change in valuation without imposing a particular functional form, we estimated non-parametric subjective weights for the seven option stimuli at each of the 24 token levels (Fig. 1d; see Methods, Non-parametric choice model). In all three monkeys, the subjective weights of earning options generally decreased, whereas those of spending options generally increased, as tokens accumulated, showing a state-dependent shift in the relative weights favoring spending over earning.

We next formalized this account using a parametric utility model based on diminishing marginal utility (see Methods, Parametric utility model). We assumed a power-law utility function *U* (*H*) = *H*^*α*^, where *α* is the curvature exponent, with marginal utility *U* ^′^(*H*) = *αH*^*α*−1^. The model integrated current token holdings *H* with the offer attributes to estimate earning and spending values.

Probability of choosing to earn was a sigmoid function of the value difference in all three monkeys (Fig. 1e and Extended Data Fig. 1c). Across 150 recording sessions (W, 26; U, 61; T, 63), the free-*α* model yielded lower held-out loss than the fixed-*α* model in all three subjects (W, *p* = 9.34 × 10^−6^; U, *p* = 1.92 × 10^−11^; T, *p* = 5.17 × 10^−12^; two-sided Wilcoxon signed-rank tests; Extended Data Fig. 1a). Independent session-wise fits characterized the distribution of fitted utility curvature across recording sessions (median *α*: W, 0.72; U, 0.81; T, 0.79; Extended Data Fig. 1b). Moreover, the value difference predicted reaction time: responses were slower near the fitted behavioral indifference point (Extended Data Fig. 1d). Together, these results support a token-holding-dependent earning-spending preference and relate the value difference between the earning and spending options to both choice and reaction time.

### 2.3 Distributed prefrontal tuning represents current token holdings

Having established that current token holdings systematically shifted earning–spending preferences, we next asked how they were represented by individual prefrontal units. A total of 2,790 units were included in the analyses: 715 in ACC, 1,113 in DLPFC and 962 in OFC (Extended Data Fig. 2). We divided the amount of tokens into five approximately logarithmically spaced groups ([1], [2–3], [4–7], [8–15] and [16–24] tokens) to accommodate the compressed scaling commonly observed for numerical magnitude [24, 25], while keeping trial counts more comparable across groups.

Units in all three regions showed significant selectivity for token holdings. Fig. 2a,b shows an OFC unit responding most strongly in the low wealth state, whereas Fig. 2c,d shows a DLPFC unit with maximal firing in the intermediate wealth state. Across ACC, DLPFC and OFC, preferred wealth levels were distributed across all five groups (Fig. 2e and Extended Data Fig. 3a).

**Fig. 2.**
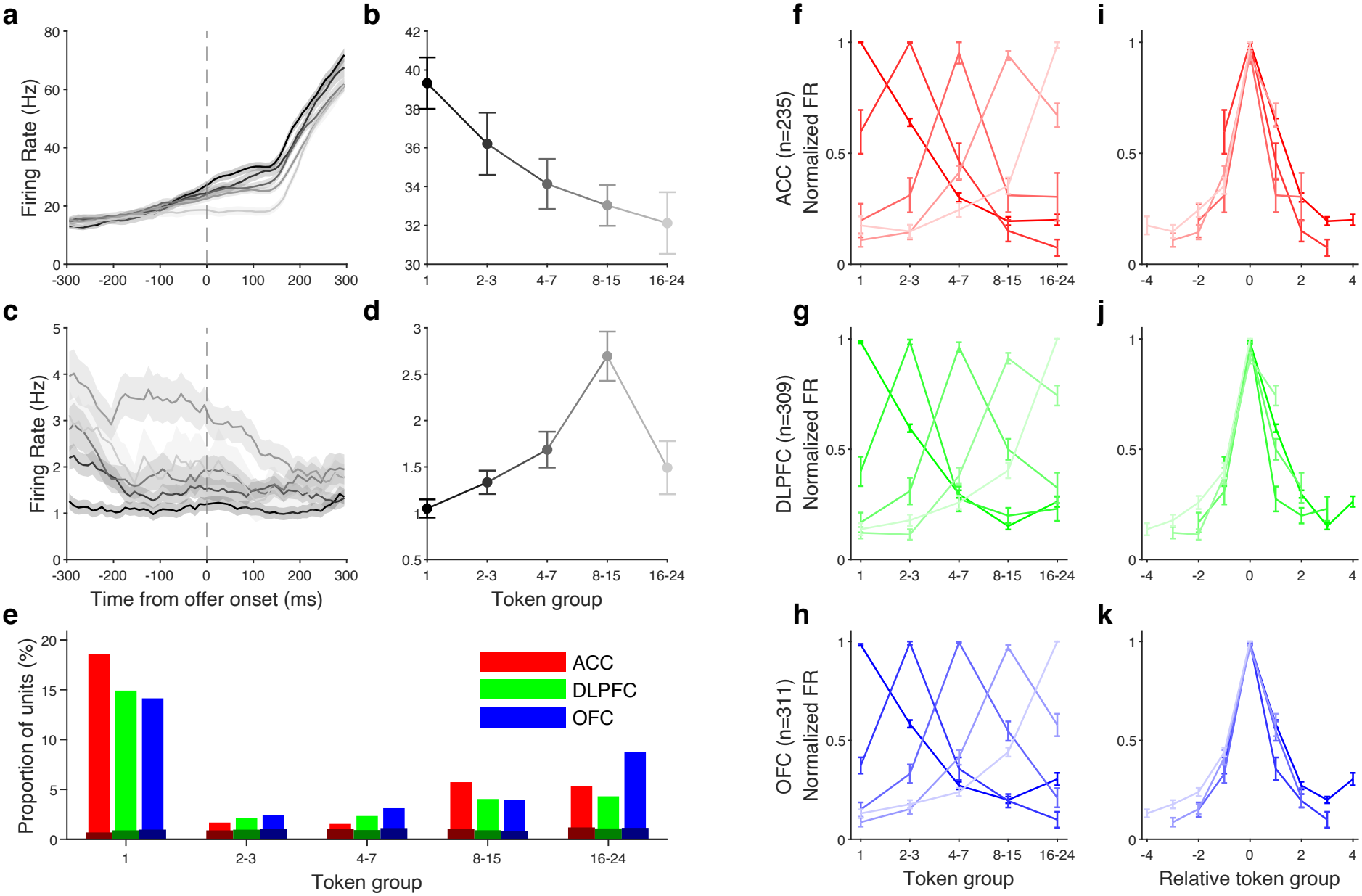
Distributed prefrontal tuning represents current token holdings. **(a–d) Example token-holding-selective units.** Peristimulus time histograms (mean ± SEM) aligned to offer onset are shown for an OFC unit (a) and a DLPFC unit (c), together with their mean firing rates in the [−300, 300) ms analysis window (b and d). Token holdings were divided into five approximately logarithmically spaced groups ([1], [2–3], [4–7], [8–15] and [16–24] tokens). Darker-to-lighter shades indicate increasing token holdings. The OFC unit responded most strongly in the [1]-token group (a,b), whereas the DLPFC unit showed maximal firing in the intermediate [8–15]-token group (c,d). **(e) Distribution of preferred token-holding groups**. Bars show the percentage of analyzed units that were selective for token holdings and preferred each group in ACC (red), DLPFC (green) and OFC (blue); dark bars show the shuffle control. **(f–h) Population tuning grouped by preferred token-holding group**. Selective units were grouped according to their preferred token-holding group. Negatively tuned curves were inverted before min– max normalization, and normalized curves were averaged within each group for ACC (f), DLPFC (g) and OFC (h). Shade denotes the preferred group from lowest (dark) to highest (light). **(i–k) Tuning re-centered on the preferred token-holding group**. The curves in f–h were shifted to express token holdings relative to each group’s preferred level for ACC (i), DLPFC (j) and OFC (k). Normalized firing rates were higher at relative distances of ±1 than at ±2 (ACC, *p* = 6.49 × 10^−31^; DLPFC, *p* = 2.61 × 10^−32^; OFC, *p* = 6.78 × 10^−40^; two-sided Wilcoxon rank-sum tests). Error bars denote SEM across trials (b,d) and across units (f–k).

We next examined the shape of the units’ tuning functions for token holdings. Units preferring different token-holding groups exhibited similarly shaped tuning curves centered at different points along the token-holding axis (Fig. 2f–h). Re-centering the curves on their preferred group revealed a common Gaussian-like profile in all three regions, with firing rates declining progressively as token holdings moved away from the preferred level (Fig. 2i–k). The well-aligned Gaussian-like tuning curve was not an artifact: normalized responses were higher at relative distances of ±1 than at ±2 from the preferred group in all three regions (ACC, *p* = 6.49 × 10^−31^; DLPFC, *p* = 2.61 × 10^−32^; OFC, *p* = 6.78 × 10^−40^; two-sided Wilcoxon rank-sum tests), a difference that was not significant after token-holding-group labels were shuffled (all *p* ≥ 0.184; Extended Data Fig. 3b). Together, tuning curves of similar shape whose preferred levels tiled the range of token holdings provided a distributed population representation of animals’ current token holdings.

### 2.3 Temporal dynamics and choice-dependent updating of token-holding representations

As the animals were constantly earning and spending tokens, the corresponding prefrontal representation of current token holdings should track their dynamics. We therefore applied cross-temporal decoding based on targeted dimensionality reduction (TDR) [26] (see Methods, Targeted dimensionality reduction). At each time bin, a TDR-derived token-holding axis was fitted independently and tested at every other bin, yielding a 24 × 24 matrix of encoding strength, quantified as squared Pearson correlations (*r*^2^) spanning the offer-onset and choice-onset epochs (Fig. 3a and Extended Data Fig. 4). Here, choice onset denotes the wallet update starting an intermediate choice within continuous fixation, with token holdings and other variables assigned to that ensuing choice (see Methods, Task events). During the offer-onset epoch, decoding was strongest near the temporal diagonal, suggesting a dynamic representation of token number. By contrast, decoding generalized broadly across the choice-onset epoch, demonstrating a stable token-holding representation during this period. Trial averaging in the cross-temporal analysis could obscure the transition structure associated with each earning and spending event. We therefore projected population activity from −50 to 150 ms relative to choice outcome (see Methods, Task events) onto TDR-derived choice and token axes (Fig. 3b). Earning drove activity toward the direction of more tokens in ACC and OFC, whereas spending drove it toward the opposite direction. To quantify these shifts, we regressed the token-holding-axis projection against time separately for earning and spending trials within each resampling iteration. The resulting temporal coefficient measured the direction and rate of movement along the token axis (Fig. 3c and Supplementary Table 1). We defined choice-dependent token updating as the joint presence of a significantly positive earning coefficient and a significantly negative spending coefficient. In the pooled data, ACC (earning, *p* = 6.98 × 10^−4^; spending, *p* = 9.55 × 10^−10^) and OFC (earning, *p* = 6.54×10^−17^; spending, *p* = 1.81×10^−4^) satisfied this bidirectional criterion, whereas DLPFC did not (earning, *p* = 8.19 × 10^−3^; spending, *p* = 1.12 × 10^−1^). Neither choice-label nor token-label shuffling yielded bidirectional updating in any region (in each case, at least one coefficient was non-significant, *p* ≥ 0.131). Thus, ACC and OFC not only represented current token number but also updated that representation when the wallet changed, providing an updated state signal that could influence subsequent value coding and decision making.

**Fig. 3.**
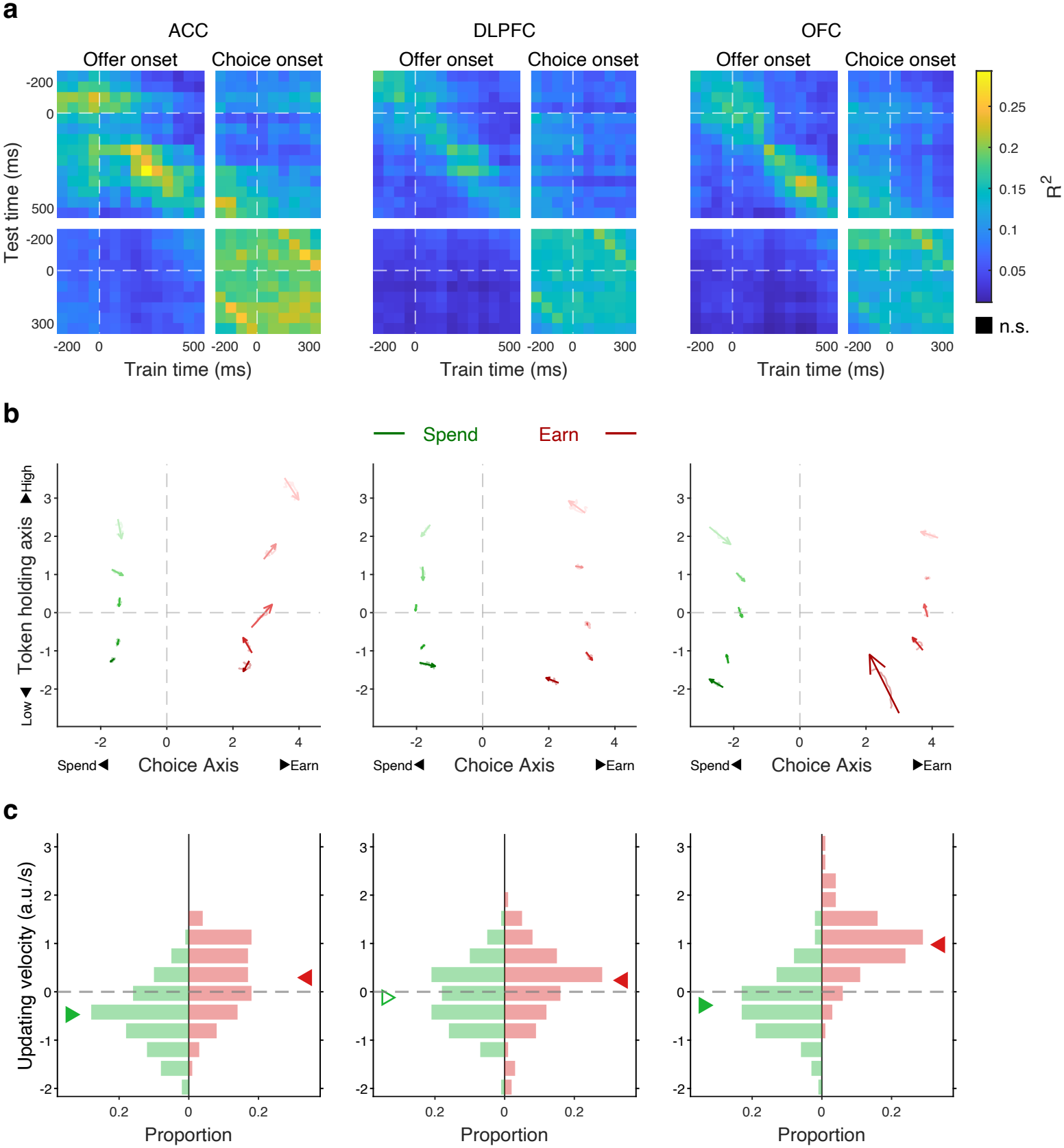
Temporal dynamics and choice-dependent updating of token-holding representations. (a) Temporal dynamics of token-holding representations. Columns show pooled population data from ACC, DLPFC and OFC. Each entry reports encoding strength, quantified as the squared Pearson correlation (*r*^2^) on held-out data, obtained using a TDR-derived token-holding axis trained at one time bin and tested at another. The four blocks comprise decoding within the offer-onset epoch (upper left; 14 bins, −200 to 500 ms relative to offer onset), within the choice-onset epoch (lower right; 10 bins, −200 to 300 ms relative to choice onset), and across the two epochs (upper right and lower left); bin width was 50 ms. Horizontal and vertical white dashed lines mark offer onset and choice onset. Black entries did not significantly exceed the shuffle control (one-sided Wilcoxon rank-sum test, *p* ≥ 0.05); no entry was black in the pooled data, so all entries exceeded the shuffle control. **(b) Population trajectories around choice outcome**. Neural population activity from −50 to 150 ms relative to choice outcome was projected onto TDR-derived choice and token-holding axes. The choice axis was oriented from spending (left) to earning (right), and the token-holding axis from lower (bottom) to higher (top) token holdings. Red and green trajectories denote earning and spending trials, respectively; darker shades denote lower token holdings. Arrows are centered on the mean trajectory positions and show the direction of the two-dimensional velocity fitted across all 20 time points; arrow lengths represent speed on a common scale across regions. **(c) Quantification of directional updating along the token-holding axis**. Histograms show the distributions of temporal coefficients (updating velocity, a.u./s) along the token-holding axis around choice outcome across resampling iterations for earning (red) and spending (green). Triangles mark medians; filled and open triangles indicate significant and non-significant shifts, respectively. Coefficients were tested against zero using two-sided Wilcoxon signed-rank tests.

### 2.4 Token holdings modulate prefrontal offer-value representations

Having established that prefrontal populations represented and updated the current token holdings, we next asked whether this state was incorporated into the neural processing of earning and spending values. The behavioral model suggested that diminishing marginal utility would reduce earning values as tokens accumulated, whereas spending values would retain the separation attributable to reward magnitude (Fig. 4a,b). Example OFC neurons exhibited response patterns consistent with the behavioral model (Extended Data Fig. 5a,b).

**Fig. 4.**
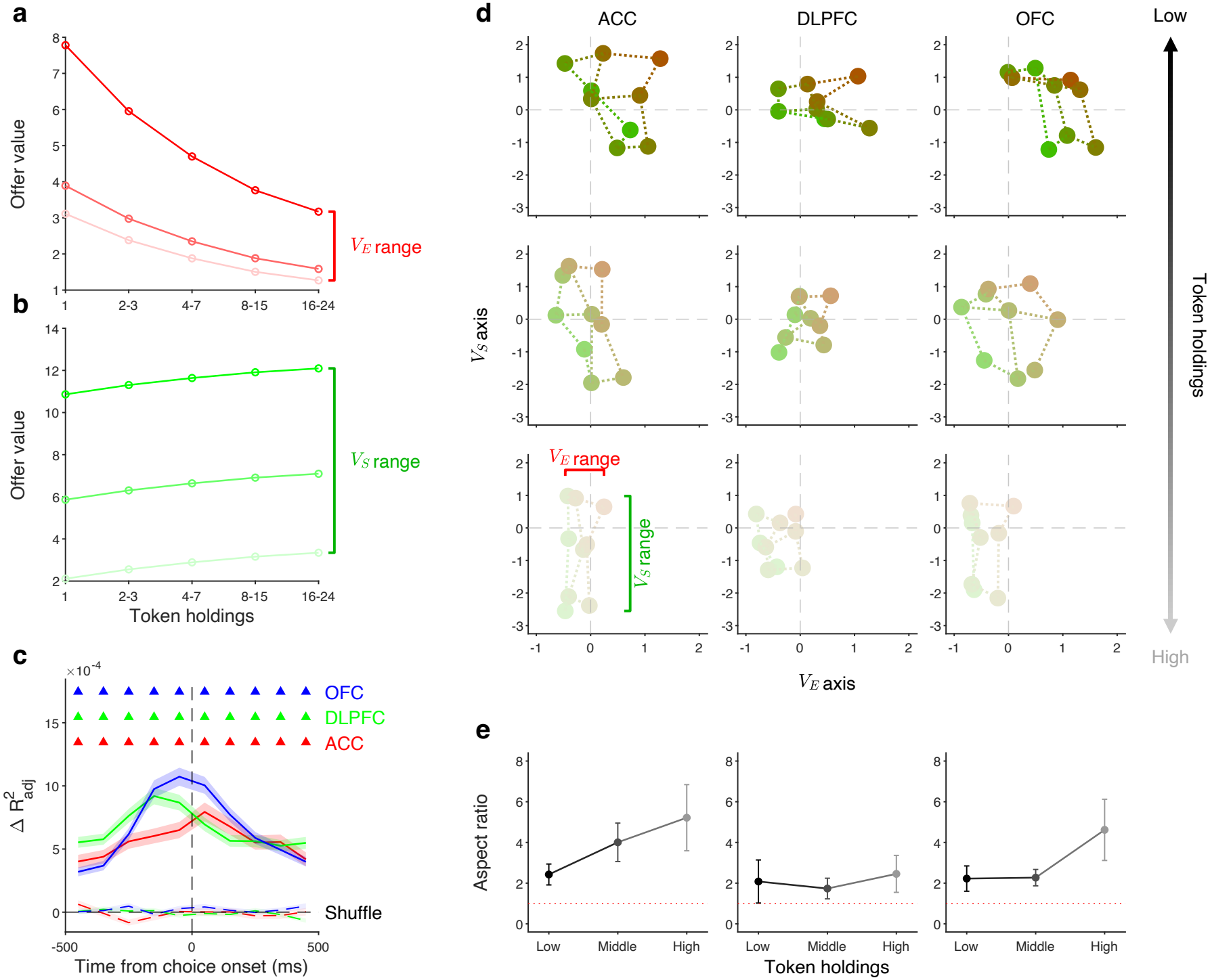
Token-holding modulation of prefrontal offer-value representations. (a–b) Behavioral-model predictions. Model-derived earning value (*V*_*E*_ ; a) and spending value (*V*_*S*_; b) are plotted across five token-holding groups ([1], [2–3], [4–7], [8–15] and [16–24] tokens). In a, the three red curves correspond to earning rates of 4, 5 and 10 tokens/s; darker red denotes a higher earning rate. In b, the three green curves correspond to spending-reward levels 1, 2 and 4 (Fig. 1b); the lowest level, 0, was rarely chosen and was excluded; darker green denotes a larger reward. Red and green brackets indicate the earning- and spending-value ranges, respectively. **(c) Incremental variance explained by token-holding-dependent earning value**. Solid lines show the across-unit mean increase in explained variance when the token-holding-dependent earning value *V*_*E*_ was added to the baseline model (*M*_1_) to form *M*_2_, aligned to choice onset; shading denotes ±SEM. Red, green and blue denote ACC, DLPFC and OFC, respectively; dashed lines show the corresponding shuffled-*V*_*E*_ controls. The vertical dashed line marks choice onset. Colored triangles denote time bins in which observed 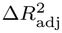 exceeded the shuffled control (*p <* 0.05, paired two-sided Wilcoxon signed-rank test across units, applied separately to each time bin without multiple-comparison correction). **(d) Population value spaces across token-holding groups**. TDR identified a two-dimensional value space with an earning-value axis (*V*_*E*_, horizontal) and a spending-value axis (*V*_*S*_, vertical). Columns show ACC, DLPFC and OFC; rows show low (1 token), middle (2–5 tokens) and high (6–24 tokens) token holdings. Each point is the condition-averaged population state for one earning–spending offer combination over [0, 150) ms after choice onset. The nine points correspond to the 3 × 3 retained subset of the behavioral condition matrix (Fig. 1b). Point hues follow the offer-condition color map in Fig. 1b, from conditions relatively favoring spending (green) to those relatively favoring earning (red). Lighter shades denote higher token holdings. Dotted lines connect neighboring conditions in the 3 × 3 offer matrix to make its geometry visible. Red and green brackets illustrate the ranges occupied along the *V*_*E*_ and *V*_*S*_ axes, respectively. One resampling iteration selected for illustration is shown (see Methods, Population value-space geometry). **(e) Aspect ratio of the population value representation**. For ACC, DLPFC and OFC, the spending-axis range was divided by the earning-axis range within each token-holding group. Points and error bars show the mean ±SD across resampling iterations; the red dotted line indicates an aspect ratio of 1 (equal spending- and earning-axis ranges).

We used a nested-model comparison to examine state-dependent value encoding beyond the independent effects of offers and holdings. Relative to a baseline model (*M*_1_) containing state-independent offer values, categorical token-holding terms and choice, a second model (*M*_2_) additionally included the token-holding-dependent earning value *V*_*E*_ derived from the behavioral model (see Methods, Nested-model comparison).

Around choice onset, the increase in explained variance, 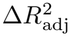, was significantly greater than the shuffled-*V*_*E*_ control in ACC, DLPFC and OFC (Fig. 4c and Extended Data Fig. 5c). This increment supports a non-additive relationship between earning-offer information and token holdings in single-unit activity around choice onset. An analogous nested-model test was not identifiable for *V*_*S*_ because its state-dependent component was additive with token holdings and was therefore absorbed by the token-holding terms already present in *M*_1_ (see Methods, Nested-model comparison).

We next used TDR analyses to examine how this state dependence was expressed in a population value space defined by earning- and spending-value axes, plotting the earning-spending conditions as an array (Fig. 4d). We quantified the geometry of this representation using the aspect ratio between the spending- and earning-axis ranges. Because the behavioral model predicted that the separation among earning values would narrow while the separation among spending values would be preserved, this ratio was expected to increase with token holdings, visualized as the relative elongation of the representational array along the *V*_*S*_ axis. The aspect ratio increased with token holdings in ACC and showed a weaker tendency in DLPFC and OFC (Fig. 4e and Extended Data Fig. 6). Token-holding-label shuffling attenuated this state-related structure, whereas value-label shuffling collapsed the underlying value-space geometry (Extended Data Fig. 6). Together, these results suggest that token holdings modulate offer-value representations across ACC, DLPFC and OFC.

### 2.5 Decision-related value representations in prefrontal cortex

The token-holding-dependent offer-value representations in prefrontal cortex point toward how the earning– spending decision may be formed. A natural decision variable is the value difference, Δ*V* = *V*_*E*_ − *V*_*S*_, but we did not observe its robust encoding at the single-unit level. A neuron encoding Δ*V* would have opposite tuning directions for *V*_*E*_ and *V*_*S*_. When we grouped neurons by their tuning directions of the two option values, we did not find more neurons had opposite tuning directions (Extended Data Fig. 7a).

An alternative is that value representations become linked to the choice, as suggested by the chosen-value neurons described in OFC [13]. In the example neuron, earning- and spending-value tuning had opposite signs within each choice condition, and both tuning directions reversed when the selected option changed (Fig. 5a,b). This pattern was consistent with encoding the value advantage of the selected option over its alternative. In the pooled populations, *V*_*E*_ encoding was stronger on earning than on spending trials (ACC, *p* = 3.69 × 10^−5^; DLPFC, *p* = 4.47 × 10^−12^; OFC, *p* = 3.90 × 10^−18^) and *V*_*S*_ encoding was stronger on spending than on earning trials (ACC, *p* = 7.33 × 10^−18^; DLPFC, *p* = 4.01 × 10^−2^; OFC, *p* = 3.90 × 10^−18^; two-sided Wilcoxon signed-rank tests across 100 resampling iterations; Fig. 5c and Supplementary Table 2). Neither choice-label nor value-label shuffling reproduced these differences in the observed direction.

**Fig. 5.**
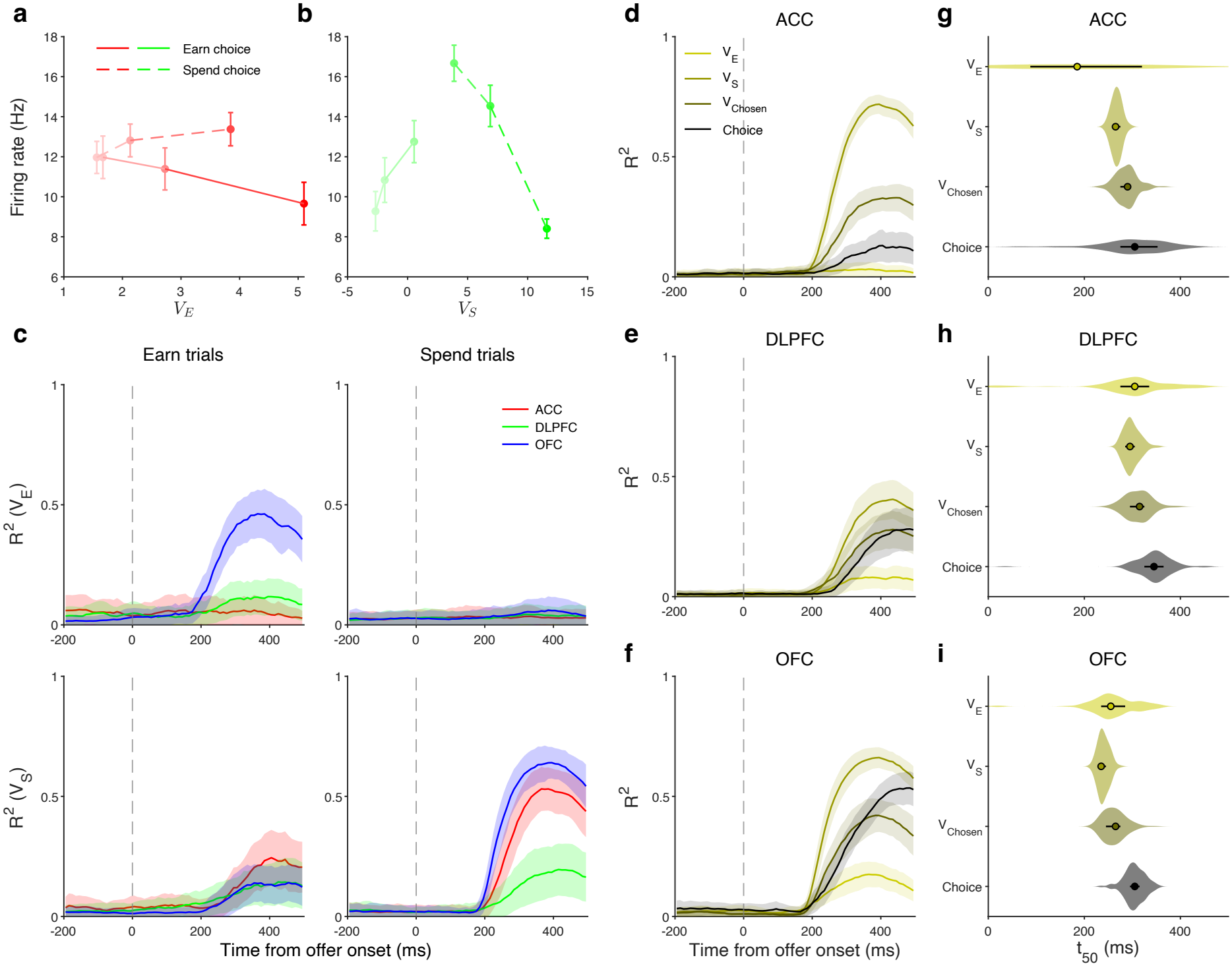
Decision-related value representations in prefrontal cortex. (a–b) Example OFC unit with choice-dependent value tuning. Mean firing rate over [0, 500) ms after offer onset is plotted against tertiles of earning value (*V*_*E*_ ; a) and spending value (*V*_*S*_; b), separately for earning (solid lines) and spending (dashed lines) choices. Red and green denote *V*_*E*_ and *V*_*S*_, respectively; error bars denote SEM. **(c) Choice-conditioned population value encoding**. TDR was applied separately to earning and spending trials. Rows show *V*_*E*_ and *V*_*S*_ encoding and columns show the selected option. Red, green and blue denote ACC, DLPFC and OFC; shading denotes ±SD across 100 iterations. **(d–f) Population encoding dynamics**. Encoding strength, measured as squared Pearson correlation (*r*^2^(*t*)), is shown for *V*_*E*_, *V*_*S*_, *V*_Chosen_ and *Choice*_ES_ in ACC (d), DLPFC (e) and OFC (f); shading denotes ±SD across 100 iterations. Light yellow, olive, dark olive and black denote *V*_*E*_, *V*_*S*_, *V*_Chosen_ and *Choice*_ES_, respectively; the same colors are used in g–i. **(g–i) Half-peak latency**. For each trajectory, *t*_50_ was the first time between offer onset and the response peak at which *r*^2^ reached the midpoint between the pre-offer baseline median and the peak. Violin area is proportional to the percentage of iterations yielding a valid estimate; points and horizontal lines denote the median and interquartile range. Between-region half-peak and onset comparisons for the value variables are reported in Supplementary Tables 7 and 8, respectively.

To capture this relationship between value and choice, we defined a relative chosen-value measure, *V*_Chosen_ = *Choice*_ES_·(Δ*V* −*b*), where *b* is the fitted choice bias and *Choice*_ES_ denotes earning (+1) or spending (−1) choice. This measure expresses the advantage of the selected option relative to the alternative and can be negative. *V*_Chosen_ was encoded at both the single-unit (Extended Data Fig. 7b) and population levels (Fig. 5d–f). The pooled population encoding strength followed *V*_*S*_ *> V*_Chosen_ *> V*_*E*_ in all three regions (Supplementary Table 3). The strengths of these representations also differed between regions. OFC represented all three value signals robustly, whereas ACC showed little *V*_*E*_ encoding during offer evaluation and DLPFC showed comparatively weak value encoding overall (Fig. 5d–f; Supplementary Table 4).

Finally, we examined the temporal order of the two offer values, relative chosen value and *Choice*_ES_ signals. For signals with stable latency estimates, half-peak and onset estimates were overall consistent with the temporal progression from *V*_*E*_*/V*_*S*_ to *V*_Chosen_ to *Choice*_ES_ (Fig. 5g–i and Supplementary Tables 5 and 6). This overall pattern is consistent with a transformation from offer-value representations toward choice, with *V*_Chosen_ as a potential intermediate representation.

### 2.6 Regional organization of good-based and action-based choice signals

Finally, choice in this task is eventually expressed as eye movements toward the left or right targets. Accordingly, we asked how the choice signals identified above were formed along the good-based and action-based coordinates. The example DLPFC unit in Fig. 6a primarily distinguished leftward from rightward responses, whereas the example OFC unit in Fig. 6b primarily distinguished earning from spending choices.

**Fig. 6.**
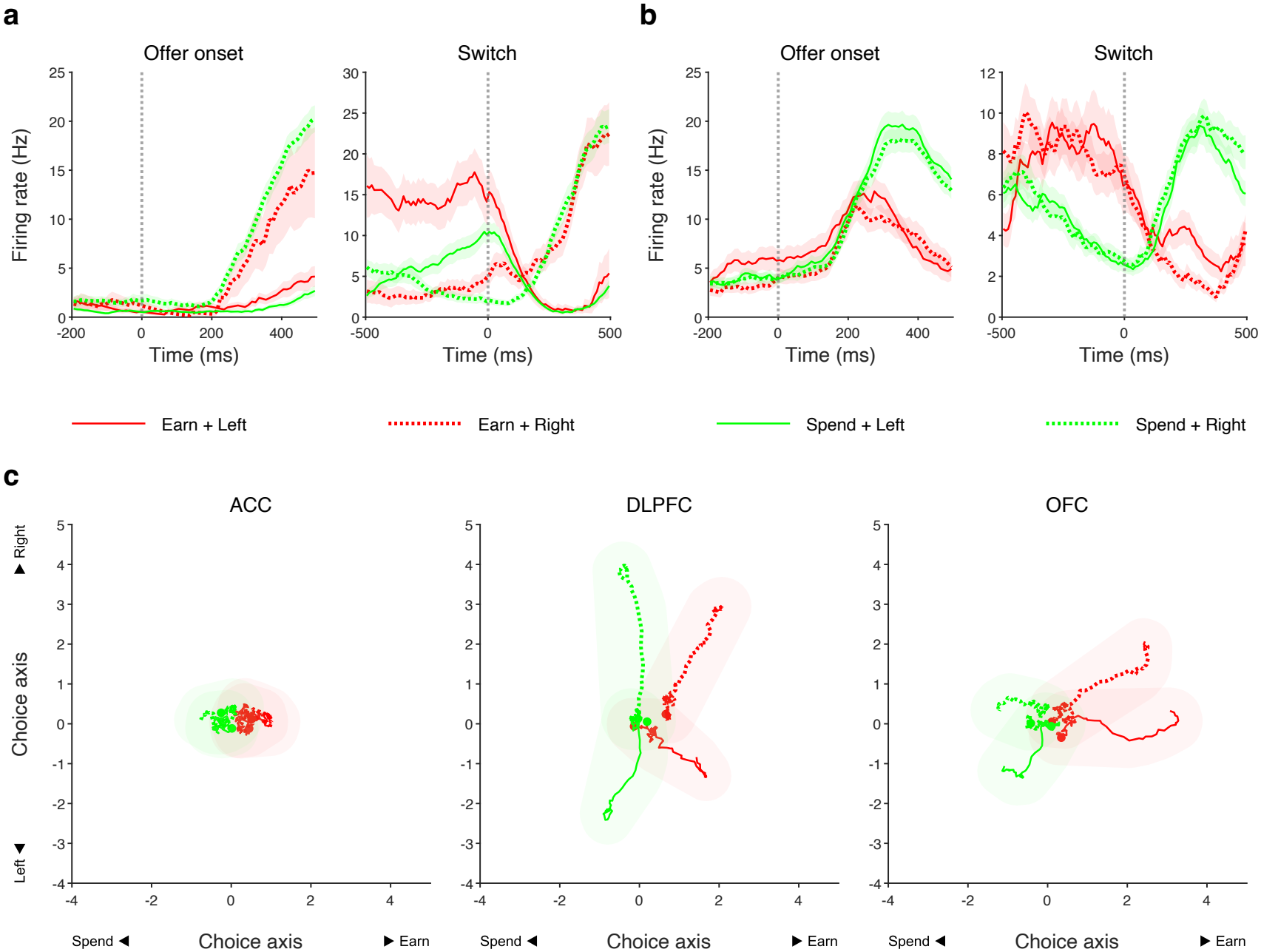
Regional organization of good-based and action-based choice signals. (a–b) Example choice-selective units. Firing rates (mean ± SEM) are shown for a DLPFC unit selective for the left/right action (a) and an OFC unit selective for the earning/spending identity of the selected option (b), aligned to offer onset (left) and choice switch (right) within each panel. Trials are grouped by earning (red) versus spending (green) choice and leftward (solid) versus rightward (dotted) response; switch-aligned trials are grouped according to the choice made after the switch. **(c) Population trajectories in good–action choice space**. TDR projected population activity onto an earning/spending choice axis (*Choice*_ES_, horizontal) and a left/right choice axis (*Choice*_LR_, vertical). Columns show ACC, DLPFC and OFC. The horizontal axis is oriented from spending (left) to earning (right). Red and green trajectories denote earning and spending choices, respectively; solid and dotted trajectories denote leftward and rightward responses. Trajectories span 10-ms bins centered from −195 to 495 ms relative to offer onset; large circles mark the population states at −195 ms. Shaded envelopes were constructed from circles centered on the mean trajectories, with radii determined by the standard deviation of pseudo-trial Euclidean distances from the mean trajectory.

We used TDR again to separate the two choice coordinates at the population level. Following offer onset, the four choice conditions separated most clearly along the earning/spending axis in OFC (Fig. 6c, right) and along the left/right axis in DLPFC (Fig. 6c, middle). The corresponding encoding trajectories and strength comparisons confirmed distinct regional biases in choice format: DLPFC preferentially encoded the left/right action, OFC preferentially encoded the earning/spending identity of the selected option, and ACC carried substantially weaker choice signals overall (Fig. 6c and Supplementary Table 9). Thus, the relative prominence of the choice representations differed by region and by choice coordinates.

We further asked whether the coordinate encoded more strongly in each region also emerged earlier. No consistent temporal advantage was evident: the relative timing of earning/spending and left/right signals varied across regions and animals and differed between onset and half-peak estimates (Supplementary Tables 10 and 11). Comparisons of each choice signal across regions likewise identified no brain area in which either signal consistently emerged first (Supplementary Tables 12 and 13). Regional preference in encoding strength was therefore not accompanied by a corresponding temporal priority.

Together, good-based and action-based choice signals were represented across the OFC and the DLPFC, with OFC preferentially encoding the earning/spending identity of the selected option and DLPFC the left/right action, but without a consistent temporal ordering across signals or regions.

## 3 Discussion

Economic decisions depend not only on the opportunities currently available but also on the resources already held. In humans, accumulated money constitutes an externally maintained resource state that preserves the capacity for future exchange and influences whether additional resources should be acquired or existing resources consumed. Here, rhesus monkeys earned virtual tokens, retained them across trials and spent them for water. As holdings increased, monkeys shifted systematically from earning toward spending, consistent with a state-dependent utility model in which the relative value of further acquisition declined while the relative value of consumption increased. Neural populations in ACC, OFC and DLPFC represented current holdings, and ACC and OFC updated these representations following resource gains and expenditures. Moreover, resource state modulated option-value coding and reshaped its population geometry across all three regions. Thus, accumulated tokens were represented not merely as previous outcomes, but as a persistent decision context linking past choices to the valuation of future opportunities.

A central contribution of this study is the characterization of an external resource state. Previous studies of state-dependent valuation have focused largely on internal physiological states, such as hunger, thirst and satiety [7, 27, 28]. These states alter the current desirability of primary rewards; for example, selective satiety reduces both behavioral preference and OFC responses to the consumed reward [9, 29]. Token holdings differ in several important respects. They were external to the body, discretely quantifiable and able to increase or decrease without a corresponding change in immediate primary-reward consumption. Whereas satiety reflects the organism’s current physiological relation to a reward, token holdings reflect the externally stored resources available for obtaining that reward.

Internal and external states nevertheless pose a common computational problem. Both must be integrated with current opportunities to determine context-appropriate values. Hunger can increase the value of acquiring food, whereas abundant external resources can reduce the marginal value of further acquisition and increase the attractiveness of consumption. Our results therefore extend the principle of state-dependent valuation from internal motivational conditions to an acquired, externally maintained resource state. Importantly, these states can change independently or even in opposite directions. Earning increased token holdings without producing immediate water, whereas spending decreased holdings while delivering water and potentially increasing satiety. The present findings build on, but are distinct from, earlier studies of token accumulation and reward saving [20–23, 30]. In many token tasks, however, holdings indicate progress toward a fixed threshold that triggers a reset and automatic delivery of a primary reward [21, 23]. Here, tokens instead functioned as spendable secondary resources: animals could voluntarily acquire, preserve or exchange them at intermediate levels. Holdings represented a continuously changing set of future possibilities rather than simply progress toward a predetermined outcome. The principal advance is consequently not only the observation that neurons encode token number, but also the identification of distributed mechanisms that represent and update spendable holdings and use them to modulate acquisition and consumption decisions.

Current holdings were represented across ACC, OFC and DLPFC, paralleling the distributed representation of internal physiological states such as hunger, thirst and satiety [11, 31, 32]. Individual neurons preferred different levels of holdings, collectively spanning the resource-state space. Such diversity in preferred levels could support a population representation that distinguishes intermediate resource levels and accommodates nonlinear relationships between holdings and behavior. Its distributed nature further suggests that resource information is available to multiple stages of decision making, including state monitoring, valuation and action selection.

Nevertheless, the three regions did not appear to use this information identically. In particular, ACC and OFC showed bidirectional updating following changes in holdings: earning shifted population activity toward states associated with higher holdings, whereas spending shifted it toward lower holdings. This result is important because it suggests that neural activity tracked the current balance of the resource account rather than merely cumulative reward history, task progression or elapsed time. Such nonspecific signals would not necessarily reverse following expenditure. DLPFC represented current holdings but did not show equally robust bidirectional updating, suggesting that representing a resource state and updating it may be partly separable operations. ACC and OFC may contribute more directly to revising the resource account after outcomes, whereas DLPFC may read out the updated state during subsequent choice and action preparation. Because recordings were not simultaneous, however, the present data cannot establish where updating originates or how resource information flows between regions.

Resource state was not simply represented in parallel with option values. It changed how those values were encoded. State-dependent value variables explained neural activity beyond models containing state-independent offer attributes, categorical holdings and choice. At the population level, increasing holdings compressed the neural separation among earning values relative to that among spending values, reshaping value geometry across all three regions. This result distinguishes representing resource state from using resource state. If holdings and option values were encoded independently, changes in holdings might translate population activity while preserving relationships among option values. Instead, holdings altered those relationships, indicating that resource state modulated the relative neural scaling of acquisition and consumption opportunities.

This mechanism provides a potential parallel with internal-state-dependent valuation. Satiety changes neural sensitivity to food-related outcomes [9, 33]; similarly, token holdings changed neural sensitivity to earning and spending opportunities. In both cases, state may reconfigure value representations so that neural distances reflect the behavioral relevance of options under current conditions. The critical difference lies in the origin of the modulating signal: satiety is an internal physiological state, whereas holdings are acquired external resources whose value derives from future exchange. Our findings therefore suggest that prefrontal circuits can apply state-dependent valuation principles to both internal needs and externally maintained economic resources.

Together, the findings support a recurrent account of economic choice: current holdings provide a resource state; this state modulates earning and spending values; valuation guides choice; and the outcome updates the resource state for the next decision. Resource state is thus both an outcome of previous behavior and an input into future valuation. This closed loop captures a defining feature of economic behavior that is absent from tasks in which decisions are independent and reward delivery terminates the relevant sequence.

## 4 Methods

### 4.1 Experimental procedures

#### 4.1.1 Animals and ethics

Three male rhesus monkeys (*Macaca mulatta*; monkeys W, U and T), aged 3–4 years at the start of the experiment, were used. Their body weights during the experiments were 6.3–8.0, 3.5–4.6 and 5.8–6.7 kg, respectively. Daily fluid intake was regulated to maintain task motivation. All procedures followed the protocol approved by the Animal Care Committee of Shanghai Institutes for Biological Sciences, Chinese Academy of Sciences (CEBSIT-2021004).

#### 4.1.2 Experimental setup and behavioral task

Monkeys sat in a custom primate chair with the head fixed and viewed a 23.6-inch monitor at a distance of approximately 60 cm. Visual stimulus presentation and reward delivery were controlled using MonkeyLogic 2 running in MATLAB. Eye position was sampled at 1,000 Hz using an EyeLink 1000 infrared eye tracker. Water was delivered through a computer-controlled solenoid valve calibrated at approximately 0.2 *µ*l ms^−1^.

Monkeys performed an earning–spending task (Fig. 1a,b) in which tokens accumulated across trials and could subsequently be spent for water. Current holdings were displayed by a central “wallet”, a 5×5 circle array approximately 4.3^°^ wide. Filled circles indicated the number of tokens currently held. The wallet remained visible during the inter-trial interval (ITI), when the option stimuli were absent, and updated immediately after each successful earning or spending event. Holdings were bounded between 0 and 25 tokens. Earning at 25 tokens did not add another token, and spending at 0 tokens did not remove one, although the attempted choice was recorded. Each session began at 0 tokens.

Each trial presented one earning stimulus and one spending stimulus. Each option stimulus was approximately 2^°^ in diameter. Immediately before option presentation, the monkey’s gaze position relative to the screen center was measured. The two options appeared 6^°^ from the screen center, one 90^°^ clockwise and the other 90^°^ counterclockwise from this gaze direction. Assignment of earning and spending stimuli to the two locations was randomized across trials. The earning rate and spending reward were sampled independently with equal probability, yielding 12 offer conditions. The three earning rates were one token per 250, 200 or 100 ms of continuous fixation (4, 5 or 10 tokens/s). Spending occurred once per 200 ms (5 tokens/s). Each spending event deducted one token and provided the water amount specified by the spending stimulus. Water amounts per token were approximately 0, 4, 7 and 11 *µ*l for monkeys W and U, and 3, 5, 7 and 14 *µ*l for monkey T.

Monkeys selected an option by directing gaze to it and maintaining fixation within a 3^°^-diameter window centered on the option. Continued fixation generated multiple token updates, and monkeys could switch between the two options before trial end. Trial duration was 1,000 ms plus an exponentially distributed interval with a mean of 2,000 ms, redrawn whenever the total exceeded 5,000 ms; the ITI was approximately 1 s. The reward magnitudes above were fixed throughout all recording sessions.

#### 4.1.3 Surgery, anatomical localization and neural recordings

Before training, each monkey was implanted with a headpost; after behavior stabilized, a recording chamber (25 mm anterior–posterior × 20 mm medial–lateral) was implanted over prefrontal cortex (left hemisphere in W and T, right in U), and the bone within it was removed with the dura left intact. Surgeries were performed under aseptic conditions and general anesthesia, induced with ketamine hydrochloride (5–15 mg kg^−1^, i.m.) and maintained with 1.5–2% isoflurane, with continuous physiological monitoring.

Recording sites were localized using post-implantation structural MRI (Siemens MAGNETOM Prisma, 3 T) and physiological landmarks encountered during electrode penetrations. ACC recordings corresponded approximately to BA 24, OFC to BA 11 and DLPFC to BA 46 (Extended Data Fig. 2).

Neuronal activity was recorded using 16-channel Plexon S-Probes (Plexon Inc., Dallas, TX, USA). Adjacent recording contacts were separated by 100 *µ*m, such that the 16 contacts spanned 1.5 mm along the probe shaft. Probes were advanced to the target depth using an EPS multichannel microdrive (Alpha Omega Engineering, Nof HaGalil, Israel). Signals were acquired using an AlphaLab SnR system (Alpha Omega Engineering) at 44 kHz and band-pass filtered online between 300 and 6,000 Hz. No task-responsiveness criterion was applied during offline unit inclusion. Electrophysiological data were collected over 150 recording sessions (26 in monkey W, 61 in U and 63 in T). After unit inclusion (see Spike sorting and unit inclusion), the dataset comprised 715 units from ACC (251 from monkey W and 464 from T; no ACC recordings were obtained from U), 1,113 from DLPFC (110, 481 and 522 from W, U and T) and 962 from OFC (106, 435 and 421 from W, U and T).

### 4.2 Data preparation

#### 4.2.1 Task events

We define *offer onset* as the simultaneous appearance of the two option stimuli, an *initial choice* as the first option fixation that successfully produced an earning or spending event, and a *wallet update* as an individual token addition or deduction. Within a period of continuous fixation, a *choice* is the interval between two consecutive wallet updates. The wallet update ending choice *k* is its *choice outcome* and simultaneously the *choice onset* of choice *k* + 1. A *choice switch* is a transition to the other option in which both the preceding and subsequent fixations produced a token event and the inter-fixation interval was nonnegative and shorter than 150 ms.

#### 4.2.2 Event inclusion

All behavioral sessions in which electrophysiological data were recorded were included. Behavioral analyses used only the first choice on each trial, provided that it was successfully executed, thereby avoiding dependence introduced by repeated within-trial selections. Observations at the wallet boundaries (0 or 25 tokens) were also excluded because one option could not change the token holdings. Consequently, all behavioral value models were evaluated over holdings of 1–24 tokens.

Neural analyses aligned to within-fixation wallet updates included only intermediate choices, excluding the first and last choice of each continuous fixation, so that no saccade occurred at the aligned wallet updates.

#### 4.2.3 Task variables and notation

Token holdings, denoted by *H*, are the number of tokens displayed in the wallet at the alignment event specified for each analysis. Choice-onset analyses assigned each event the earning/spending choice beginning at that update and the post-update token holdings; choice-outcome analyses assigned the choice ending at that update and the pre-update token holdings. Value predictors were computed from the offer information and token holdings assigned in this way. In neural models, *H* entered as a continuous predictor (1–24 tokens), as a categorical predictor with five token-holding groups, or as a categorical predictor with one level per token count (up to 24 levels), as stated for each model. The five token-holding groups were [1], [2–3], [4–7], [8– 15] and [16–24] tokens. These groups were approximately logarithmic, yielding more comparable trial counts across groups. Geometry analyses used three approximately logarithmic groups—1, 2–5 and 6–24 tokens—to maintain sufficient events per offer-condition cell.

*V*_*E*_ and *V*_*S*_ denote earning and spending values from the free-*α* utility model; 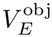 and 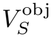 denote the state-independent predictors from the fixed-*α* model (see Parametric utility model). Δ*V* = *V*_*E*_ − *V*_*S*_ denotes the unnormalized value difference. *Choice*_ES_ denotes earning (+1) versus spending (−1), and *Choice*_LR_ denotes left (+1) versus right (−1) choices, defined by whether the chosen option appeared to the left or right of the screen center; the two options were always placed on opposite sides of the center (see Experimental setup and behavioral task). Unless otherwise stated, “choice” refers to the earning/spending choice; *Choice*_LR_ is used only in the analyses of good-based and action-based choice signals (Fig. 6). *V*_*L*_ and *V*_*R*_ are obtained by assigning *V*_*E*_ and *V*_*S*_ to the corresponding option locations. *V*_Chosen_ = *Choice*_ES_ · (Δ*V* − *b*) denotes the relative chosen-value measure, where *b* is the choice bias of the parametric utility model.

#### 4.2.4 Spike sorting and unit inclusion

Spike sorting was performed automatically using Kilosort4 [34], with each recording fragment sorted independently; units classified as good or multi-unit activity (MUA) were retained. To exclude silent periods caused by electrode drift, loss of isolation or cell injury, firing rate was evaluated in complete non-overlapping 60-s windows. For each unit, only the longest continuous sequence of windows with a mean firing rate strictly above 1 Hz was retained. Trials were required to lie completely within this active interval. At least 500 trials were required per unit.

#### 4.2.5 Firing-rate estimation

Spike trains were initially converted to firing rates in 10-ms bins. Analyses based on a single mean firing rate within a predefined window used unsmoothed rates unless otherwise stated. PSTHs and other time-resolved trajectories were smoothed with a causal Gaussian kernel (*σ* = 100 ms). The kernel spanned only [−300, 0] ms relative to each time point, so the smoothed rate depended on activity in the current and preceding bins.

### 4.3 Behavioral value models and validation

#### 4.3.1 Non-parametric choice model

To quantify the modulation of option value by token holdings (Fig. 1d) without assuming a specific functional form, we estimated subjective weights for the seven option stimuli (three earning, four spending) at each of the 24 token-holding levels. Earning probability was modeled by a sigmoid function:

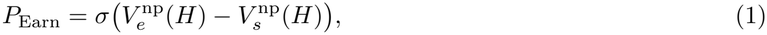

where *σ*(*x*) = 1*/*(1 + exp(−*x*)), 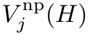 is the estimated weight of option stimulus *j* (*j* = 1, …, 7) at token holding *H*, and *e* and *s* index the earning and spending stimuli offered on that trial. To regularize the solution and encourage gradual variation across adjacent levels, we augmented the cross-entropy loss with an *L*_2_ smoothing penalty:

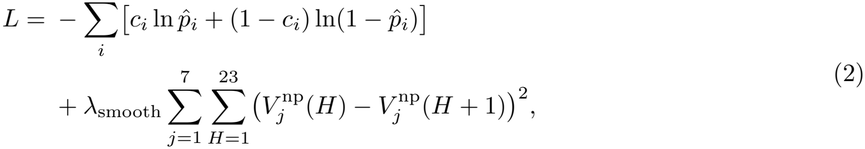

where the sum runs over trials *i*, the included trials of a monkey pooled across its sessions, *c*_*i*_ ∈ {0, 1} is the binary choice (1 = earn, 0 = spend), 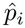 is the predicted earning probability, and *λ*_smooth_ = 10 controls the smoothing strength.

The model contained no separate bias term or fixed reference option. Choice probabilities depend on differences between option weights; a common constant offset leaves both the likelihood and the smoothing penalty unchanged. Interpretation therefore concerns relative weights and their state dependence rather than an absolute value baseline.

#### 4.3.2 Parametric utility model

We developed a parametric model based on marginal utility theory (Fig. 1e and Fig. 4a,b). We assumed a power-law utility function *U* (*H*) = *H*^*α*^, where *α* is the curvature exponent, with marginal utility *U* ^′^(*H*) = *αH*^*α*−1^. Reward magnitude was expressed in dimensionless units by dividing the water amounts specified under Experimental setup and behavioral task by the second-smallest amount within each monkey. The subjective values of earning and spending were defined as:

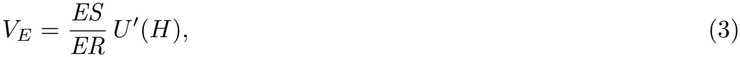

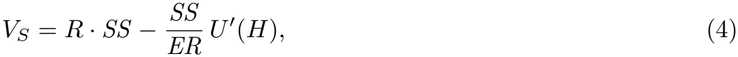

where *ES* is the earning speed (tokens/s), *R* is the normalized reward magnitude (in arbitrary units), *ER* is the exchange rate (tokens per unit reward), and *SS* = 5 tokens/s is the spend speed. The first term in *V*_*S*_ represents the immediate reward benefit; the second captures the opportunity cost of spending a token scaled by its marginal utility. Choice probability was given by:

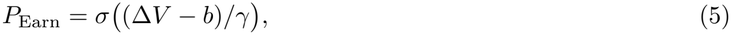

where *γ >* 0 is a temperature parameter (smaller values imply greater sensitivity) and *b* is a choice bias. Free parameters (*α, ER, b, γ*) were estimated by minimizing the cross-entropy loss, with *α* ≥ 0, *ER >* 0 and *γ >* 0; *b* was unconstrained.

The *fixed-α* model served as a nested baseline in which *α* was constrained to 1, corresponding to linear utility and constant marginal utility. All remaining parameters were estimated freely, testing whether state dependence improved prediction within this model family.

#### 4.3.3 Model validation and session-wise fitting

To evaluate out-of-sample predictive performance (Extended Data Fig. 1a), we applied leave-one-session-out (LOSO) cross-validation separately within each subject. In each fold, model parameters were estimated from the pooled data from all remaining sessions and evaluated on the held-out session. Performance on the heldout session was quantified as the average negative log-likelihood per trial (NLL per trial). The per-fold loss difference was defined as Δ*L* = *L*_fixed_ − *L*_free_, such that positive values indicate better generalization of the free-*α* model. Loss differences were tested against zero across held-out sessions using two-sided Wilcoxon signed-rank tests.

To characterize the distribution of fitted utility curvature across recording sessions (Extended Data Fig. 1b), the free-*α* model was fitted independently to each session.

#### 4.3.4 Reaction-time analysis

Reaction time (Extended Data Fig. 1d) was defined as the interval from offer onset to target acquisition (gaze entry into the fixation window of the initially chosen option). Analyses included all recording sessions and used valid initial choices at holdings of 1–24 tokens. Decision difficulty was indexed by the absolute normalized value difference, 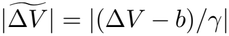, with smaller values indicating proximity to the fitted behavioral indifference point. Within each subject, trials were divided at the median into hard (at or below the median) and easy (above the median) groups. After grouping, reaction times exceeding 700 ms were excluded. Reaction-time distributions were compared within each subject using two-sided Wilcoxon rank-sum tests on pooled trials.

### 4.4 Targeted dimensionality reduction

#### 4.4.1 TDR rationale

Individual prefrontal neurons can encode mixtures of correlated task variables. Following Mante et al. [26], we used targeted dimensionality reduction (TDR) to estimate population axes that separately account for response variance associated with each variable. To avoid overestimating encoding through overfitting, axes were estimated on training events and evaluated on held-out events. Each of 100 independent resampling iterations comprised five steps: (i) split: each unit’s events were divided into training and held-out sets; (ii) regression: each unit’s training responses were regressed on the task variables; (iii) orthogonalization: coefficient vectors were stacked across units and orthogonalized by QR decomposition; (iv) readout: held-out responses were combined into pseudo-population trials and projected onto the resulting axes; and (v) quantification: encoding strength was computed from these projections. Analysis-specific response windows, predictors and their QR order, groups and group labels are listed in Table 1.

**Table 1.** TDR analysis specifications. Predictors are listed in QR column order; extracted axes are in bold; *H* is continuous (1–24 tokens). Windows are in ms relative to the stated event: [*a*:*b*:*c*] denotes consecutive *b*-ms bins from *a* to *c*, each with its own fitted axis; [*a, b*) denotes a single fitting window, over which training responses were averaged to estimate the axes. Held-out responses were projected onto these fixed axes at each time bin, except in the geometry analysis, which used window-averaged responses. ^*†*^Causally smoothed responses; other responses were unsmoothed. Token-holding groups are defined under Task variables and notation. For *r*^2^, group labels were the lower bounds of the token-holding groups (1, 2, 4, 8 and 16), the mean held-out value of each tertile averaged across units, or +1/−1 for earning/spending or left/right choices. Pseudo-trials were drawn with replacement within groups. ED, Extended Data.

| Analysis | Fitting window | Predictors (QR order) | Groups |
|---|---|---|---|
| Cross-temporal<br>Fig. 3a<br>ED Fig. 4<br>§4.5.2 | Offer onset<br>[−200:50:500]<br>Choice onset<br>[−200:50:300] | [*V*<sub>*E*</sub>, *V*<sub>*S*</sub>, *Choice*<sub>ES</sub>, ***H***] | 5 token-holding groups<br>(60 pseudo-trials per group) |
| Updating<br>Fig. 3b,c<br>§4.5.3 | Choice outcome<br>[−50, 150)<sup>†</sup> | [*V*<sub>*E*</sub>, *V*<sub>*S*</sub>, ***Choice***<sub>**ES**</sub>, ***H***] | 5 token-holding × 2 choice groups<br>(mean of 30 pseudo-trials per group) |
| Geometry<br>Fig. 4d,e<br>ED Fig. 6<br>§4.6.2 | Choice onset<br>[0, 150) | [*Choice*<sub>ES</sub>, *H*, ***V***<sub>***E***</sub>, ***V***<sub>***S***</sub>] | 3 token-holding × 9 offer groups<br>(unit-wise mean of all held-out events per group) |
| Choice-conditioned<br>Fig. 5c<br>§4.7.2 | Offer onset<br>[0, 500)<sup>†</sup> | [*H*, ***V***<sub>***E***</sub>, ***V***<sub>***S***</sub>]<br>Fitted within each choice | Value tertiles within each choice<br>(100 pseudo-trials per group) |
| Value and choice<br>Fig. 5d–i<br>§4.7.3 | Offer onset<br>[0, 500)<sup>†</sup> | [*H*, *Choice*<sub>ES</sub>, *V*<sub>*S*</sub>, *V*<sub>Chosen</sub>, ***V***<sub>***E***</sub>]<br>[*H*, *Choice*<sub>ES</sub>, *V*<sub>*E*</sub>, *V*<sub>Chosen</sub>, ***V***<sub>***S***</sub>]<br>[*H*, *Choice*<sub>ES</sub>, *V*<sub>*E*</sub>, *V*<sub>*S*</sub>, ***V***<sub>**Chosen**</sub>] | Value tertiles<br>(100 pseudo-trials per group) |
|  |  | [*H*, *V*<sub>*E*</sub>, *V*<sub>*S*</sub>, *V*<sub>Chosen</sub>, ***Choice***<sub>**ES**</sub>] | 2 choice groups<br>(150 pseudo-trials per group) |
| Choice format<br>Fig. 6c<br>§4.8 | Offer onset<br>[0, 500)<sup>†</sup> | [*V*<sub>*L*</sub>, *V*<sub>*R*</sub>, *V*<sub>*E*</sub>, *V*<sub>*S*</sub>, *H*, ***Choice***<sub>**LR**</sub>, ***Choice***<sub>**ES**</sub>]<br>Ridge, λ<sub>ridge</sub> = 1 | 2 choice groups per axis<br>(150 pseudo-trials per group) |

#### 4.4.2 Axis estimation and held-out readout

i. **Split**. Each unit’s events were split equally at random into training and held-out sets. Units were eligible if they had at least 50 valid events and at least five held-out events in each group (Table 1).
ii. **Regression**. Responses **y**_*n*_(*t*) were z-scored at each time point using training means and standard deviations, giving **z**_*n*_(*t*); the same transformation was applied to held-out responses. In all TDR models, *H* entered as a continuous predictor (1–24 tokens). Predictors were z-scored using training events so that coefficients were comparable across predictors. For unit *n*,

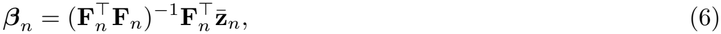

where **F**_*n*_ is the standardized training predictor matrix and 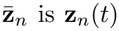 of training events averaged over the fitting window. The choice-format analysis (Fig. 6c) used ridge regression, adding 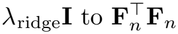.
iii. **Orthogonalization**. Eligible units were sampled without replacement, up to 300 for pooled populations and 100 for individual monkeys. Their coefficient vectors were stacked and orthogonalized by QR decomposition:

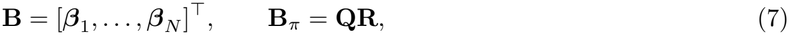

where *π* is the analysis-specific column order. The axis **q**_*v*_ is the column of **Q** corresponding to variable *v*, that is, the normalized component of the coefficient column ***β***_*v*_ orthogonal to the columns preceding it in *π*. Column orders and extracted axes are listed in Table 1.
iv. **Readout**. For held-out group *g*, pseudo-trial *m* combined one held-out event per unit, drawn with replacement from that unit’s events in *g*:

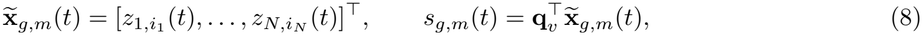

where *z*_*n,i*_(*t*) is the z-scored held-out response of unit *n* on event *i*.
v. **Quantification**. Encoding strength was quantified as

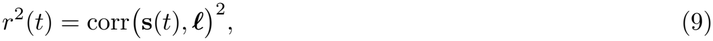

where **s**(*t*) contains the pseudo-trial projections and ***l*** their group labels.

#### Exceptions and controls

Updating (Fig. 3b,c; §4.5.3) and geometry (Fig. 4d,e; §4.6.2) instead used one mean projection per group. Shuffle controls permuted labels within units and repeated steps (i)–(v).

#### 4.4.3 Latency

Latency was measured from each *r*^2^(*t*) trajectory using the pre-offer baseline [−200, 0) ms. Let *θ* be the 95th percentile of *r*^2^(*t*) within the baseline. Onset latency was

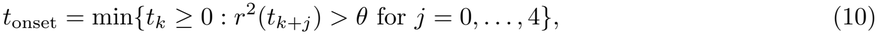

(10) where *t*_*k*_ are consecutive time bins. Half-peak time was

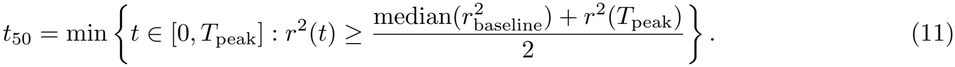

An iteration yielded no *t*_50_ if its peak did not exceed *θ*, and no onset latency if no five consecutive bins exceeded *θ*; each signal’s valid estimates were retained separately.

### 4.5 Token-holding representation and updating

#### 4.5.1 Single-unit token-holding selectivity and tuning

To quantify token-holding selectivity (Fig. 2e–k and Extended Data Fig. 3), we used OLS regression to model the mean firing rate over [−300, 300) ms relative to offer onset against task variables:

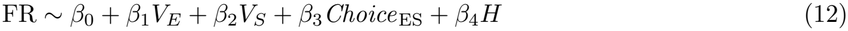

where *H* was the five-level categorical token-holding predictor at offer onset defined under Task variables and notation. Units were classified as token-holding selective if the *F* -test for the *H* term reached significance (*p <* 0.05). The preferred token-holding level was defined as the level yielding the smallest *p*-value across two-sided Wilcoxon rank-sum tests, each comparing the mean firing rate of one token-holding group against all remaining groups combined. The direction of tuning was determined by the sign of 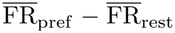. For population averaging, the tuning curves of negatively tuned units 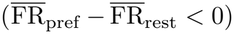 were inverted prior to normalization. Min-max normalization was then applied:

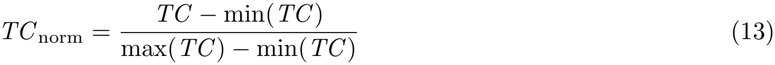

Normalized curves were averaged across units sharing the same preferred token-holding level to yield group-level population tuning curves. To characterize the shape of the re-centered population tuning curves (Fig. 2i–k), the normalized firing rates at relative positions ±1 and ±2 from each unit’s preferred token-holding level were extracted and compared using a two-sided Wilcoxon rank-sum test.

For the shuffle control, token-holding-group labels were independently permuted across trials within each unit before repeating unit selection, preferred-group assignment, sign alignment, normalization and peak recentering. Selective proportions (Fig. 2e and Extended Data Fig. 3a) were averaged across ten permutations per unit, whereas the population tuning curves and their associated statistical comparisons (Extended Data Fig. 3b) used the first permutation. Individual-monkey analyses repeated the complete procedure used for the pooled results.

#### 4.5.2 Cross-temporal decoding

For cross-temporal decoding (Fig. 3a and Extended Data Fig. 4), at each 50-ms training bin, a token-holding axis was fitted using *V*_*E*_, *V*_*S*_, *Choice*_ES_ and continuous *H* (1–24 tokens) and applied to held-out responses at all bins in both epochs, yielding a 24 × 24 matrix. Encoding strength was averaged across iterations.

For the shuffle control, token-holding labels (*H*) were permuted across events within each unit, separately for each epoch; *V*_*E*_, *V*_*S*_ and *Choice*_ES_ were left unchanged. Each matrix entry was compared with the shuffle distribution using a one-sided Wilcoxon rank-sum test; entries not exceeding shuffle at *p <* 0.05 were displayed in black.

#### 4.5.3 Choice-dependent updating of token-holding representations

For choice-dependent updating (Fig. 3b,c and Supplementary Table 1), TDR models with *V*_*E*_, *V*_*S*_, *Choice*_ES_ and continuous *H*, in this QR order, were fitted to responses averaged over [−50, 150) ms relative to choice outcome. Held-out projections onto the *Choice*_ES_ and *H* axes were obtained for the five token-holding groups crossed with earning and spending (Table 1). Axes were oriented toward earning and higher token holdings using pre-update group means. Within each iteration, each token-group trajectory was referenced to its first sample and regressed on time; the temporal coefficient, defined as the mean of the five group slopes, was computed separately for earning and spending updates; positive and negative coefficients indicated movement toward higher and lower represented holdings, respectively. The displayed trajectories used an independent split and all eligible units.

Temporal coefficients were tested against zero using two-sided Wilcoxon signed-rank tests. Bidirectional updating required a significantly positive earning coefficient and a significantly negative spending coefficient (both *p <* 0.05). Choice-label and token-holding-label shuffles permuted *Choice*_ES_ or *H*, respectively, across events within each unit, with the other predictors unchanged, and repeated regression, QR and projection; pooled analyses included both controls, and real-data analyses were repeated for individual monkeys.

### 4.6 Token-holding-dependent offer-value encoding

#### 4.6.1 Nested-model comparison

For nested-model comparisons (Fig. 4c and Extended Data Fig. 5c), 10-ms firing rates were smoothed with the causal Gaussian kernel described above and averaged in ten non-overlapping 100-ms windows spanning −500 to 500 ms relative to choice onset. Units required at least 100 valid events spanning all five token-holding groups.

Two nested linear models were fitted in each window. The baseline model (*M*_1_) included state-independent offer values, choice and token holdings:

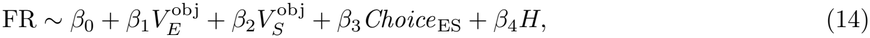

where 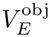 and 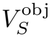 are token-holding-independent values estimated from equations (3) and (4) with *α* = 1. *H* entered as a categorical predictor with one level per token count (1–24), so that any effect of holdings alone, however nonlinear, was absorbed by the baseline. The token-holding-modulated model (*M*_2_) added the free-*α* earning value:

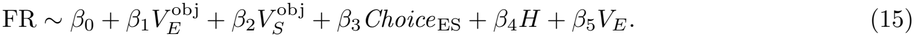

Because *V*_*E*_ is a product of an offer term and a holdings term (equation (3)), it is not spanned by *M*_1_. An analogous increment could not be estimated for *V*_*S*_: as shown in equation (4), *V*_*S*_ is an additive combination of an offer term and a holdings term, both already spanned by 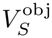 and the categorical *H* in *M*_1_. Model performance was quantified by adjusted *R*^2^ to penalize the additional parameter in *M*_2_. The modulation index 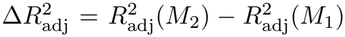 was computed for every unit and time window. For the shuffled-*V*_*E*_ control, the added token-holding-dependent earning-value predictor was permuted once across all valid events within each unit; firing rates and baseline predictors were unchanged. Observed and shuffled per-unit increments were compared using a paired two-sided Wilcoxon signed-rank test in each 100-ms time bin.

#### 4.6.2 Population value-space geometry

We used TDR-derived *V*_*E*_ and *V*_*S*_ axes to visualize the joint population representation of earning and spending offers around choice onset (Fig. 4d,e and Extended Data Fig. 6). The task contained 12 offer conditions formed by crossing three earning rates with four spending reward magnitudes. The lowest spending-reward option was rarely chosen and therefore provided too few observations for stable condition averages. We excluded this condition from the geometry analysis, leaving the 3 × 3 subset formed by the three earning rates and the three retained spending-reward levels. The three token-holding groups were defined under Task variables and notation. For each unit, z-scored held-out responses were averaged across all events within each of the 27 conditions (nine offer conditions × three token-holding groups), yielding one condition mean per unit; these unit-wise means were stacked across units and projected onto the axes to obtain 27 population states. The TDR model included *Choice*_ES_, continuous *H, V*_*E*_ and *V*_*S*_, in this QR order, fitted to responses averaged over [0, 150) ms after choice onset; the *V*_*E*_ and *V*_*S*_ axes were extracted (Table 1).

To summarize the scale of the projected geometry along each value axis, the nine coordinates on each axis were sorted within every token-holding group and resampling iteration. We defined the trimmed earning-axis and spending-axis ranges as

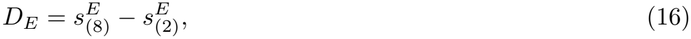

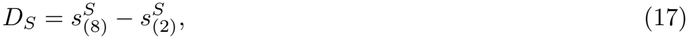

where 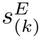 and 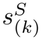denote the *k*th ordered projections on the *V*_*E*_ and *V*_*S*_ axes, respectively. Excluding the single most extreme coordinate at each end reduced the influence of any one condition mean. Within each token-holding group and resampling iteration, the aspect ratio was defined as

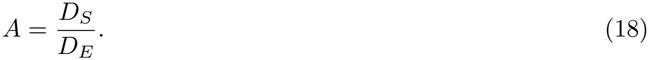

An aspect ratio greater than 1 indicates that the offer representation extends further along the spending-value axis than along the earning-value axis. Aspect ratios and component ranges were summarized as mean ± SD across all 100 resampling iterations. For display (Fig. 4d), we considered only iterations in which *D*_*E*_ changed strictly monotonically across the three token-holding groups in the direction predicted by the behavioral model (ACC, 64; DLPFC, 14; OFC, 25 of 100 iterations). Among these, the iteration maximizing the product of the eight squared Pearson correlations between condition-center projections and model values (*V*_*E*_ and *V*_*S*_, within each token-holding group and across all groups) multiplied by the magnitude of the slope of *D*_*E*_ across token-holding groups was shown.

Two shuffle controls were applied to the pooled dataset. Before the training/held-out split, either token counts (token-holding-label shuffle) or offer values (*V*_*E*_ and *V*_*S*_ as pairs; value-label shuffle) were permuted across all eligible events within each unit. The complete procedure was then repeated.

### 4.7 Decision-related value representations

#### 4.7.1 Single-unit value selectivity

##### Joint encoding of earning and spending values

To test whether single units preferentially encoded the value difference Δ*V* (Extended Data Fig. 7a), we fit an OLS regression model to mean firing rates over [0, 500) ms after offer onset and over [0, 150) ms after choice onset:

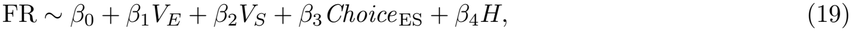

where *H* entered as a categorical predictor with one level per token count observed for the unit (up to 24 levels). Fits required at least 20 valid events per unit in the corresponding epoch. Units with a significant *β*_1_ (*p <* 0.05) were classified as 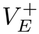 or 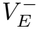 according to its sign, and units with a significant *β*_2_ as 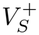 or 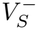. Among units significant for both *V*_*E*_ and *V*_*S*_, we constructed a 2 × 2 contingency table (rows: *V*_*E*_ sign; columns: *V*_*S*_ sign) and applied a two-sided Fisher’s exact test to assess sign association. We inspected its direction to determine whether the anti-diagonal cells (opposite-sign tuning: 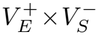 and 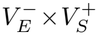) were over-represented, as would be expected if Δ*V* were the dominant encoded quantity.

##### Relative chosen-value selectivity

We defined a relative chosen-value measure *V*_Chosen_ as

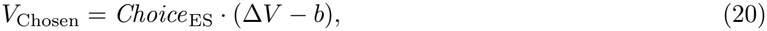

where *b* is the choice bias of the parametric utility model (see Parametric utility model). This variable is the signed value advantage of the chosen option over the unchosen option relative to the fitted indifference point. The regression model was

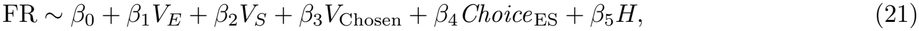

with *H* again a categorical predictor with one level per observed token count, fit to mean unsmoothed firing rates over [0, 500) ms after offer onset. Fits required at least 20 valid offer-onset events per unit. This model was used for all five encoding proportions in Extended Data Fig. 7b. Selectivity for *V*_*E*_, *V*_*S*_, *V*_Chosen_ and *Choice*_ES_ was assessed using two-sided tests of the corresponding regression coefficients (*p <* 0.05). Token-holding selectivity was assessed using a joint partial *F* -test comparing the full model with a reduced model omitting *H* (*p <* 0.05). For these five encoding proportions, empirical false-positive proportions were estimated using ten response permutations within each unit. The response vector was permuted across valid offer-onset events while the complete predictor matrix was held fixed before refitting the same model. Shuffle proportions were averaged across repetitions.

#### 4.7.2 Choice-conditioned value encoding

For choice-conditioned value encoding (Fig. 5c and Supplementary Table 2), earning and spending event pools were downsampled without replacement to equal sizes before training/test splitting. Units were eligible if both choice pools contained at least 50 events after downsampling, with at least five held-out events in each *V*_*E*_ and *V*_*S*_ tertile within each choice. Separately for earning and spending events, TDR models with continuous *H, V*_*E*_ and *V*_*S*_, in this QR order, were fitted to responses averaged over [0, 500) ms after offer onset, and held-out projections onto the *V*_*E*_ and *V*_*S*_ axes were grouped by value tertiles within each choice (Table 1).

Within each iteration, encoding strength was averaged over [0, 500) ms after offer onset. We defined 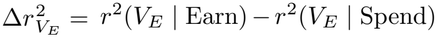 and 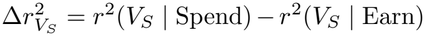 and tested each against zero using a two-sided Wilcoxon signed-rank test. For the shuffle controls, either *Choice*_ES_ labels (choice-label shuffle) or offer values (*V*_*E*_ and *V*_*S*_ as pairs; value-label shuffle) were permuted across all valid events within each unit, and the complete procedure, starting from downsampling, was then repeated. The complete procedure was repeated for individual monkeys.

#### 4.7.3 Population encoding of value and choice

For population encoding of value and choice (Fig. 5d–i and Supplementary Tables 3–8), a joint model included continuous *H* (1–24 tokens), *Choice*_ES_, *V*_*E*_, *V*_*S*_ and *V*_Chosen_ (Table 1).

Encoding strength was averaged over [0, 500) ms after offer onset (Fig. 5d–f). Strengths of *V*_*E*_, *V*_*S*_ and *V*_Chosen_ were compared between signals within each region using two-sided paired Wilcoxon signed-rank tests (Supplementary Table 3) and between regions using two-sided Wilcoxon rank-sum tests (Supplementary Table 4). Half-peak and onset latencies (Fig. 5g–i) were estimated as defined under Latency. Within each region, *V*_*E*_, *V*_*S*_ and *V*_Chosen_ were each compared with *Choice*_ES_, and *V*_*E*_ and *V*_*S*_ with *V*_Chosen_ (Supplementary Tables 5 and 6); the three value signals were also compared between regions (Supplementary Tables 7 and 8). Latency comparisons used two-sided Wilcoxon rank-sum tests on each signal’s valid estimates.

### 4.8 Good-based and action-based choice representations

The population choice-format model (Fig. 6c and Supplementary Tables 9–13) regressed responses averaged over [0, 500) ms after offer onset on *V*_*L*_, *V*_*R*_, *V*_*E*_, *V*_*S*_, continuous *H, Choice*_LR_ and *Choice*_ES_, and the *Choice*_LR_ and *Choice*_ES_ axes were extracted (Table 1). Because *V*_*L*_ + *V*_*R*_ = *V*_*E*_ + *V*_*S*_ on every event, the value predictors are exactly collinear and the OLS solution is not unique; this model was therefore fit by ridge regression with *λ*_ridge_ = 1.

Encoding strength was averaged over [0, 500) ms after offer onset. Strengths of the *Choice*_ES_ and *Choice*_LR_ signals were compared within each region using two-sided paired Wilcoxon signed-rank tests (Supplementary Table 9). Onset and half-peak latencies, defined under Latency, were compared between the two signals within each region (Supplementary Tables 10 and 11) and for each signal between regions (Supplementary Tables 12 and 13), using two-sided unpaired Wilcoxon rank-sum tests on each signal’s valid estimates. Analyses were repeated for pooled and individual-monkey populations.

The two-dimensional display used an independent split and all eligible units; each of the four groups formed by crossing earning/spending with left/right choice required at least five held-out events. Within each unit and joint group, the same 75 replacement-drawn events were projected onto both coordinates. At each time point, the standard deviation of pseudo-trial Euclidean distances from the mean trajectory defined a circle radius; an alpha shape around the combined circle points formed the displayed envelope. This envelope illustrates trajectory dispersion rather than an axis-wise standard deviation or confidence region.

## Acknowledgements

We thank Wei Kong and Lu Yu for their help in all phases of the study.

## Declarations

### Author contribution

H.W. and T.Y. designed the study. H.W. and H.S. performed the experiments and collected the data. H.W. analyzed the data. H.W., L.Z., and T.Y. wrote and revised the paper.

## Data availability

The data that support the findings of this study are available from the corresponding author upon reasonable request.

---

**Extended Data Figure 1.**
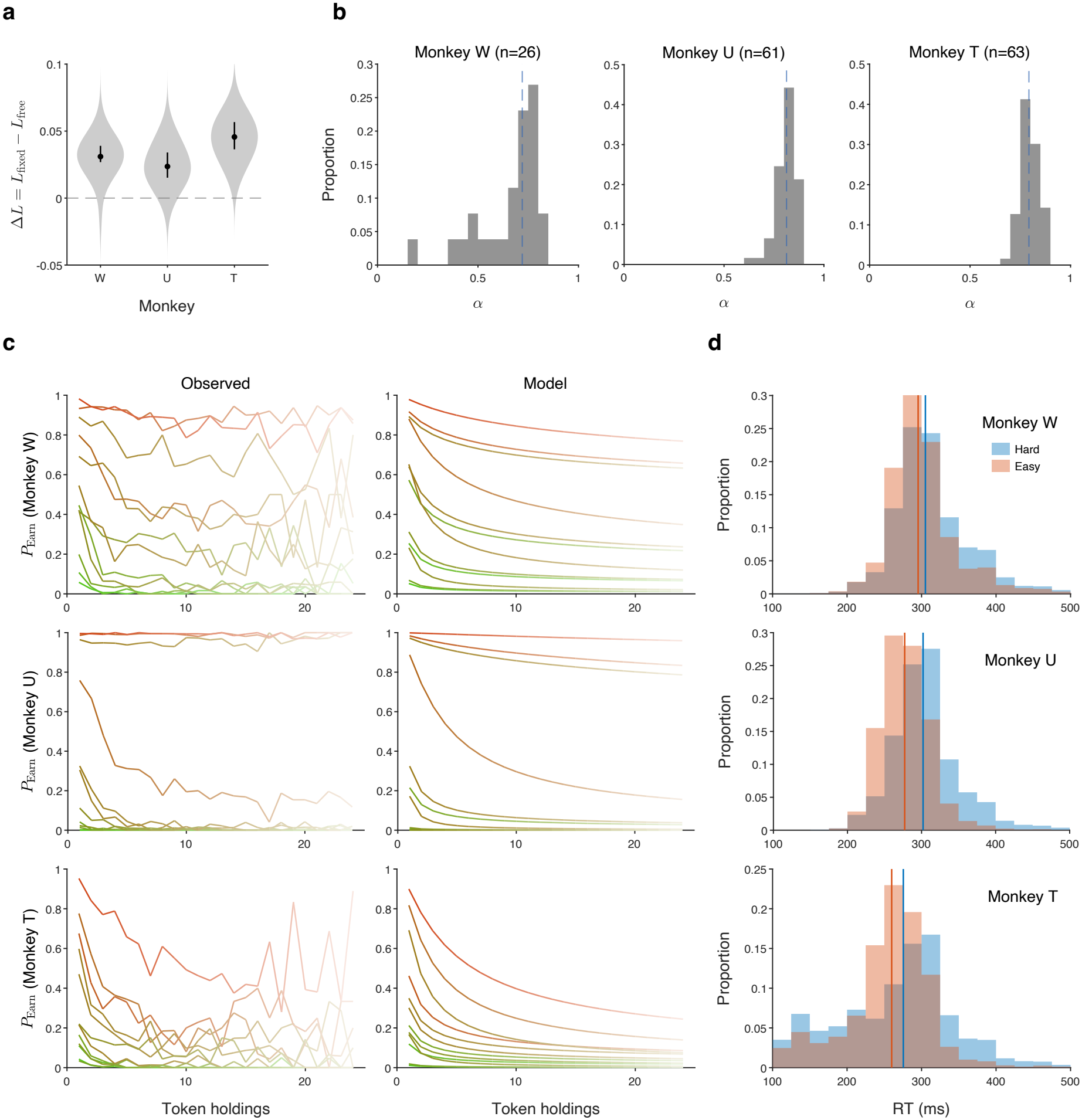
Behavioral-model validation and reaction time. **(a) Out-of-sample model comparison**. Leave-one-session-out cross-validation compared the free-*α* utility model with a nested fixed-*α* model in which *α* = 1. The ordinate shows the held-out loss difference, Δ*L* = *L*_fixed_ − *L*_free_, expressed as average negative log-likelihood per trial; positive values indicate better prediction by the free-*α* model. Violin plots show loss differences across held-out sessions, and black points and vertical lines denote the median and interquartile range, respectively. The free-*α* model had lower held-out loss in all three subjects (monkey W: *n* = 26 sessions, *p* = 9.34 × 10^−6^; monkey U: *n* = 61, *p* = 1.92 × 10^−11^; monkey T: *n* = 63, *p* = 5.17 × 10^−12^; two-sided Wilcoxon signed-rank tests against zero). **(b) Session-wise utility curvature**. To characterize variation across recording sessions, the free-*α* model was fitted independently to each session. Histograms show the proportion of sessions in each bin of the resulting *α* estimates for monkeys W, U and T (bin width, 0.05); blue dashed lines mark the medians (W, 0.72; U, 0.81; T, 0.79). **(c) Observed and model-predicted choice patterns**. Rows show monkeys W, U and T. Observed earning proportions (left) and proportions predicted by the subject-level free-*α* utility model (right) are plotted as a function of token holdings for each of the 12 offer conditions. Ordinates show earning probability (*P*_Earn_). Colors follow the offer-condition convention in Fig. 1b,c, with lighter shades indicating larger token holdings. **(d) Decision difficulty and reaction time**. Trials were divided within subject at the median absolute normalized value difference, 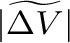, into easy (above the median; orange) and hard (at or below the median; blue) groups. Reaction time was measured from offer onset to target acquisition. Histograms show the proportion of trials in each 25-ms reaction-time bin, normalized separately for each group over 0–700 ms; vertical lines mark group medians. Trials with reaction times exceeding 700 ms were excluded after grouping; the displayed range is 100–500 ms. Responses were longer for hard than for easy trials in each subject (W, *p* = 5.50 × 10^−100^; U, *p <* 10^−300^ (numerical underflow); T, *p* = 7.53 × 10^−64^; two-sided Wilcoxon rank-sum tests on pooled trials within each subject).

**Extended Data Figure 2.**
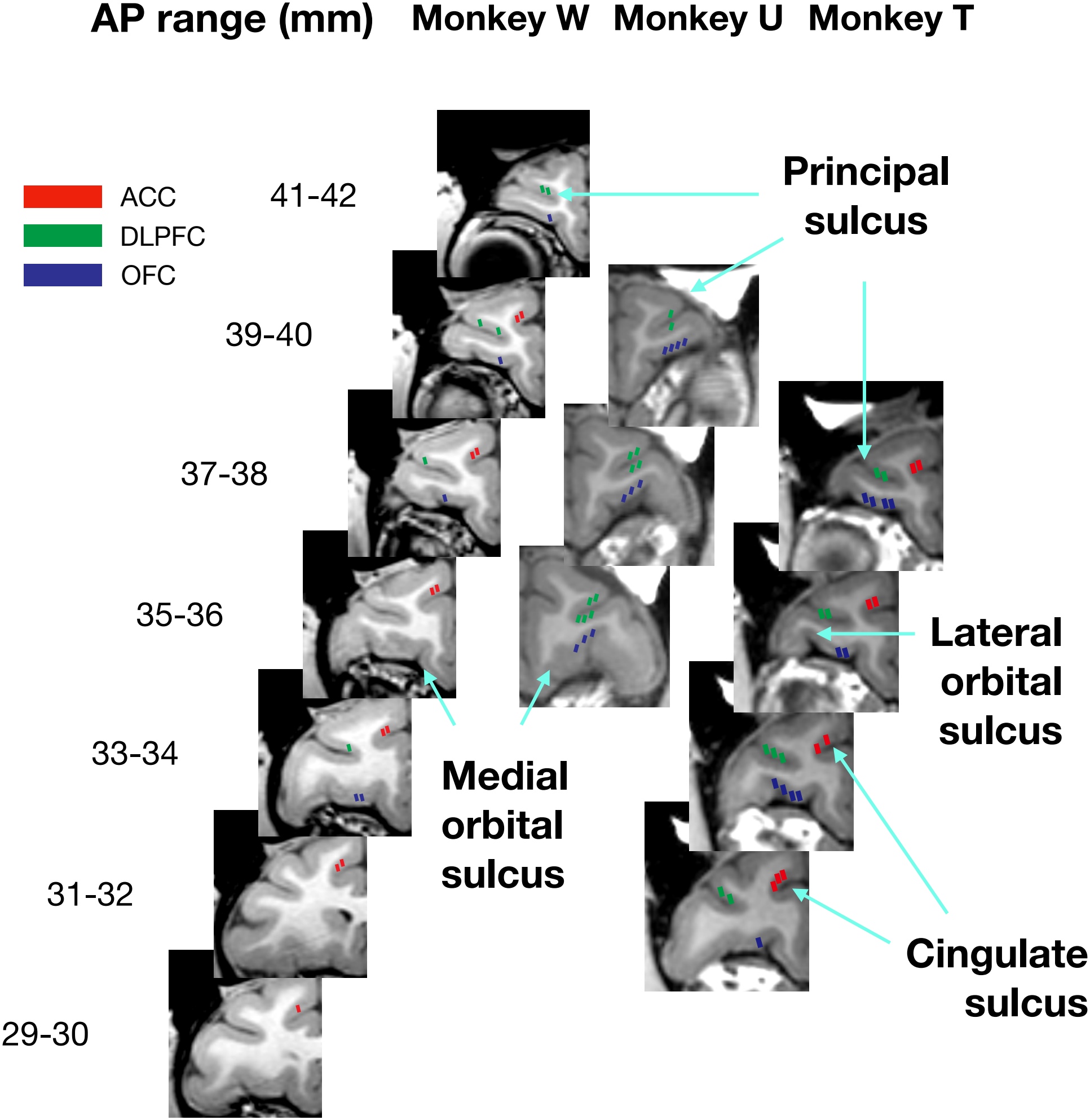
Recording-site localization. Coronal structural MR images show recording sites in monkeys W (left hemisphere), U (right hemisphere) and T (left hemisphere). Sections are arranged by anterior–posterior (AP) coordinate relative to ear-bar zero, with the AP range (mm) indicated at left. Colored rectangles mark recording sites assigned to ACC (red; approximately BA 24), DLPFC (green; approximately BA 46) and OFC (blue; approximately BA 11). Each site was sampled with a 16-channel Plexon S-Probe whose recording contacts were spaced 100 *µ*m apart and spanned 1.5 mm along the probe shaft. Rectangles are shown with a length of approximately 2 mm along the probe axis, slightly greater than the contact span, to accommodate possible electrode drift and localization uncertainty. Sites sampled in multiple sessions are shown once. Cyan arrows identify the principal, medial orbital, lateral orbital and cingulate sulci.

**Extended Data Figure 3.**
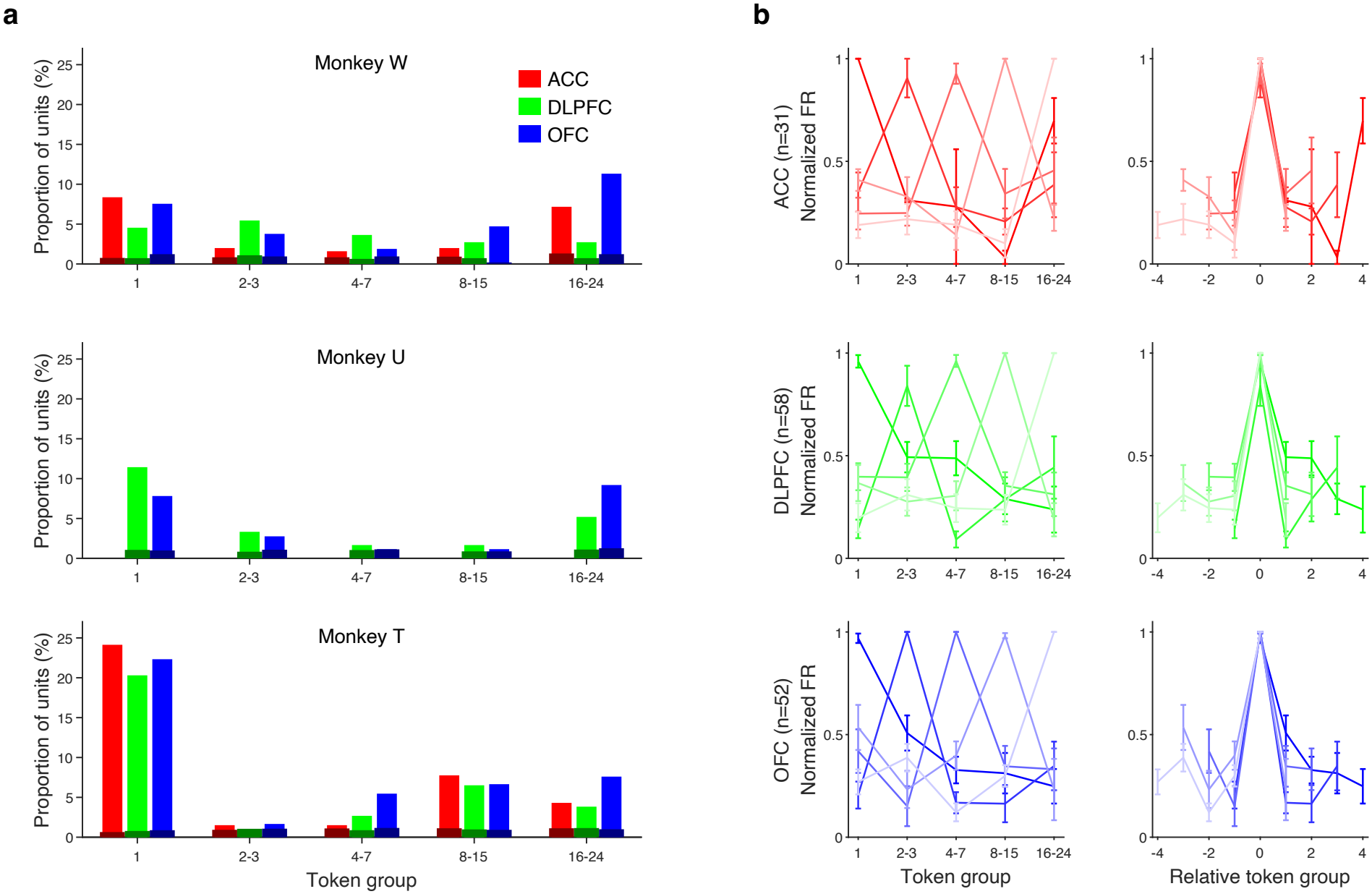
Individual-animal token-holding tuning and shuffle controls. (a) Preferred token-holding distributions in individual monkeys. Bars show the percentage of analyzed units that were selective for token holdings and preferred each group in monkeys W (top), U (middle) and T (bottom). Red, green and blue denote ACC, DLPFC and OFC, respectively; dark bars show the shuffle control. No ACC recordings were obtained from monkey U. **(b) Token-holding-label shuffle control for population tuning**. Normalized population tuning curves after token-holding-group labels were shuffled and the complete unit-selection, preferred-group assignment, normalization and peak-recentering procedure was repeated. Rows show ACC, DLPFC and OFC; left panels group curves by their preferred token-holding group, and right panels show the same curves re-centered on the preferred token-holding group. Error bars denote SEM across units. For the shuffled profiles, comparisons of normalized responses at relative distances of ±1 and ±2 were no longer significant (ACC, *p* = 0.613; DLPFC, *p* = 0.424; OFC, *p* = 0.184; two-sided Wilcoxon rank-sum tests). By contrast, the difference in the observed data remained evident in individual-animal analyses, with normalized firing rates higher at ±1 than at ±2 in every recorded region (monkey W: ACC, *p* = 7.17 × 10^−5^; DLPFC, *p* = 6.60 × 10^−3^; OFC, *p* = 7.55 × 10^−4^; monkey U: DLPFC, *p* = 4.74 × 10^−11^; OFC, *p* = 4.53 × 10^−9^; monkey T: ACC, *p* = 5.77 × 10^−28^; DLPFC, *p* = 2.02 × 10^−21^; OFC, *p* = 2.82 × 10^−30^; two-sided Wilcoxon rank-sum tests).

**Extended Data Figure 4.**
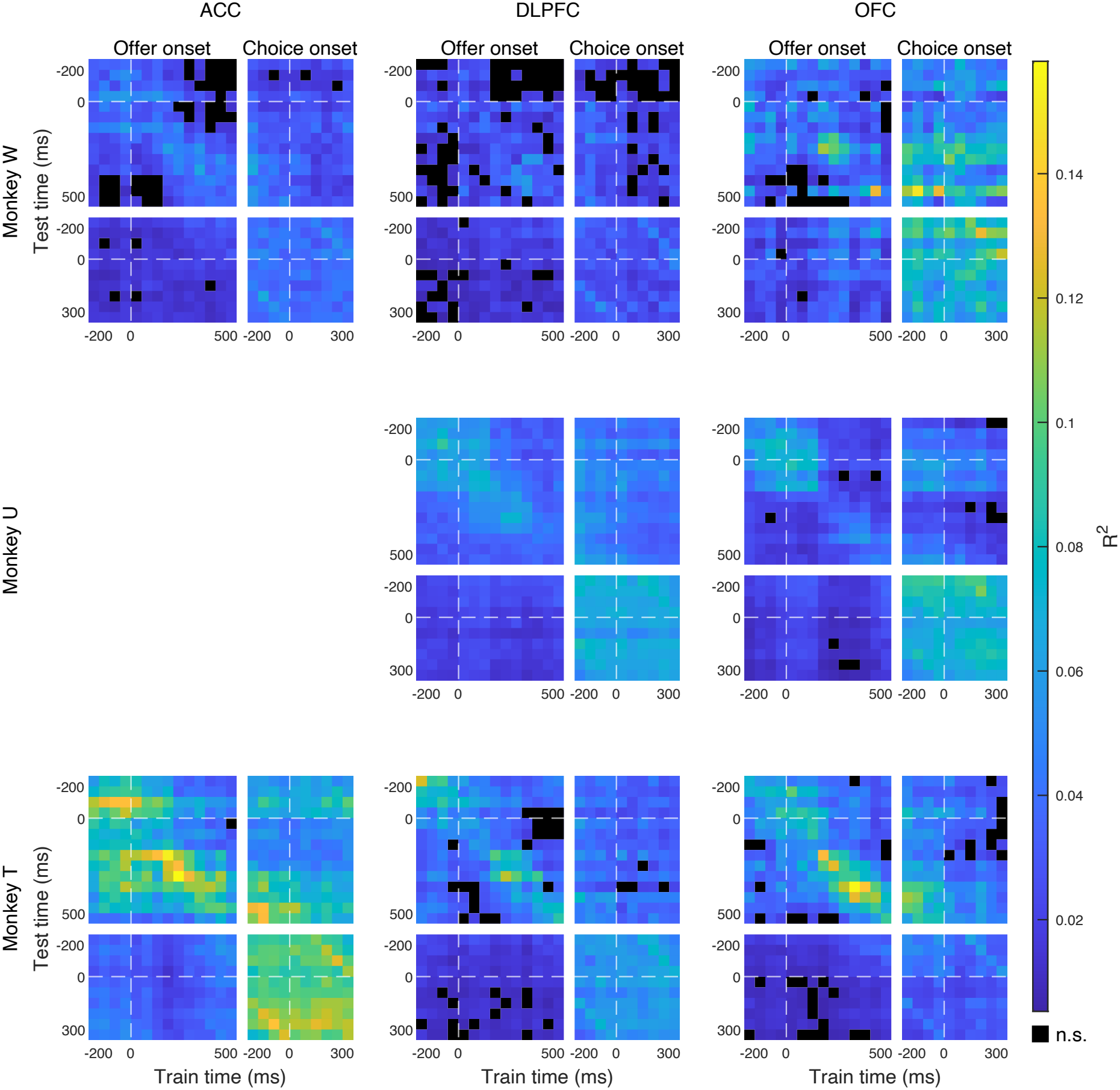
Cross-temporal decoding of token holdings in individual animals. Rows show monkeys W, U and T, and columns show ACC, DLPFC and OFC; ACC was not recorded in monkey U. Within each animal–region panel, the upper-left matrix shows offer-onset-aligned decoding (14 50-ms bins, −200 to 500 ms) the lower-right matrix shows choice-onset-aligned decoding (10 50-ms bins, −200 to 300 ms), and the off-diagonal matrices show decoding across the two epochs. Each entry is the mean squared Pearson correlation (*r*^2^) on held-out data obtained by fitting a TDR-derived token-holding axis at the training time and applying it at the test time. Horizontal and vertical dashed lines mark the corresponding event onset. Black entries did not significantly exceed the token-holding-label-shuffled control (one-sided Wilcoxon rank-sum tests comparing real and shuffled iterations, *p* ≥ 0.05).

**Extended Data Figure 5.**
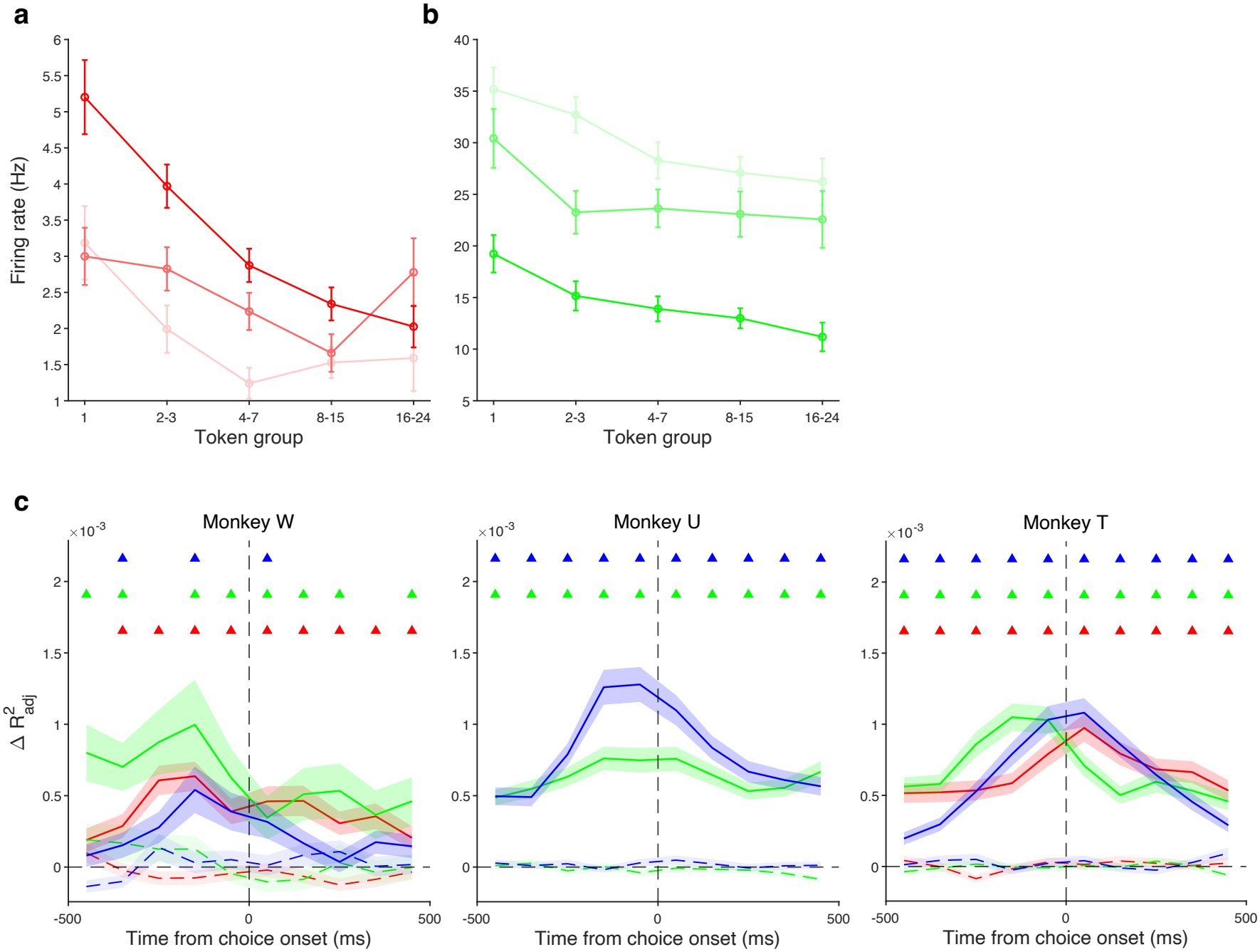
Single-unit evidence for token-holding modulation of offer-value encoding. (a–b) Example OFC units. Mean firing rates over [0, 150) ms after choice onset are plotted across the five token-holding groups for the three earning rates (a; red) or the three spending-reward levels retained for analysis (b; green); darker shades denote higher offer values and error bars denote SEM. The spending example showed negative value tuning. **(c) Token-holding-dependent earning-value model comparison in individual animals**. Columns show monkeys W, U and T. Solid lines show the across-unit mean increase in explained variance 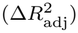 when the token-holding-dependent earning value *V*_*E*_ was added to the baseline model (*M*_1_) to form *M*_2_, aligned to choice onset; shading denotes ±SEM. Red, green and blue denote ACC, DLPFC and OFC, respectively; dashed lines show the corresponding shuffled-*V*_*E*_ controls. The vertical dashed line marks choice onset. Colored triangles denote time bins in which observed 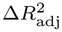 exceeded the shuffled control (*p <* 0.05, paired two-sided Wilcoxon signed-rank test across units, applied separately to each time bin without multiple-comparison correction). ACC was not recorded in monkey U.

**Extended Data Figure 6.**
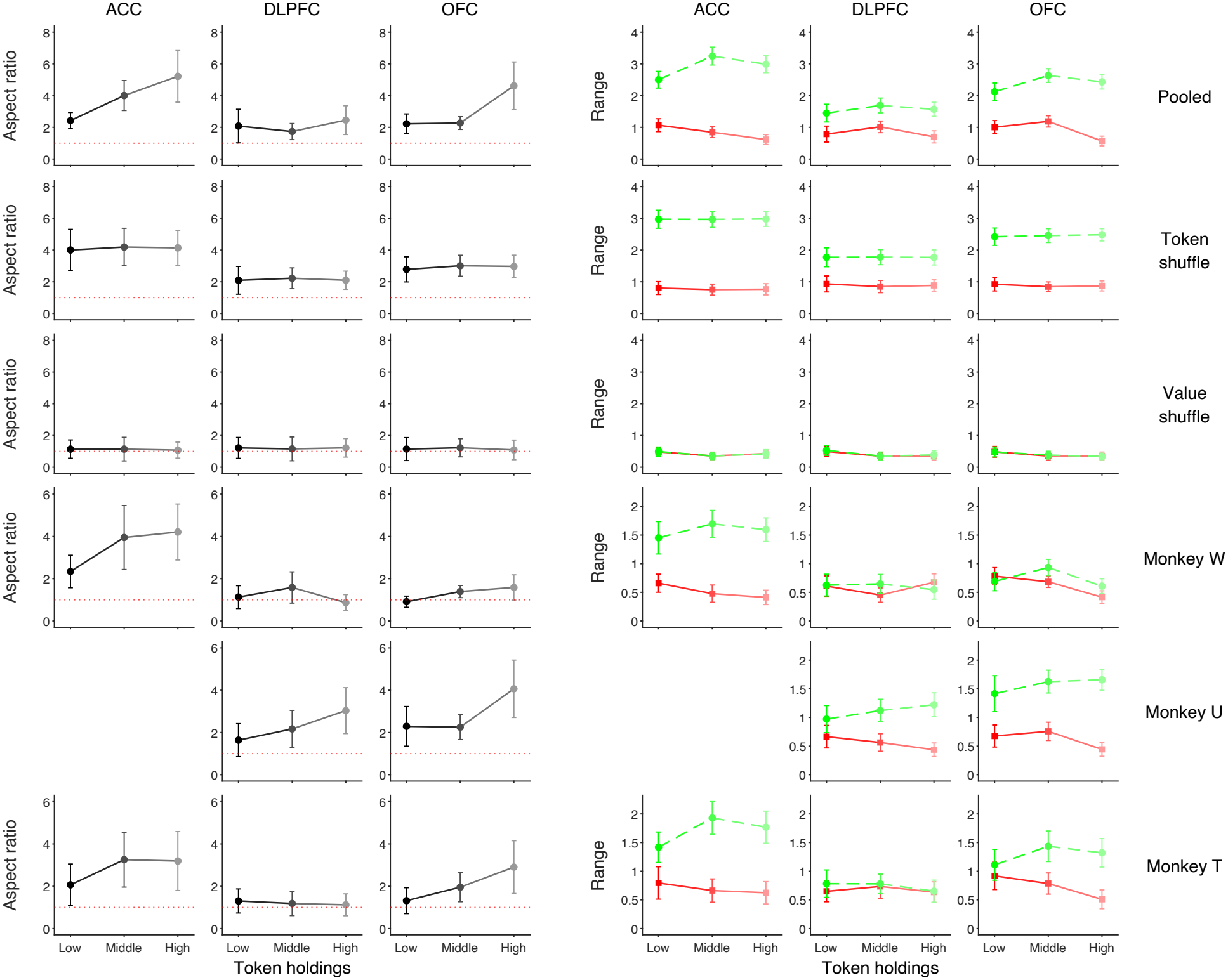
Components, controls and individual-animal results for the geometry of the population value representation. The three columns on the left show the aspect ratio (spending-axis range divided by earning-axis range) for ACC, DLPFC and OFC; the three columns on the right show its component earning-axis (red solid) and spending-axis (green dashed) ranges. Points and error bars show the mean ±SD across resampling iterations. Red dotted lines indicate an aspect ratio of 1. Rows show the pooled population, the complete analysis after shuffling token-holding labels, the complete analysis after shuffling value labels, and the corresponding results for monkeys W, U and T. ACC panels are blank for monkey U because no ACC recordings were obtained from this animal.

**Extended Data Figure 7.**
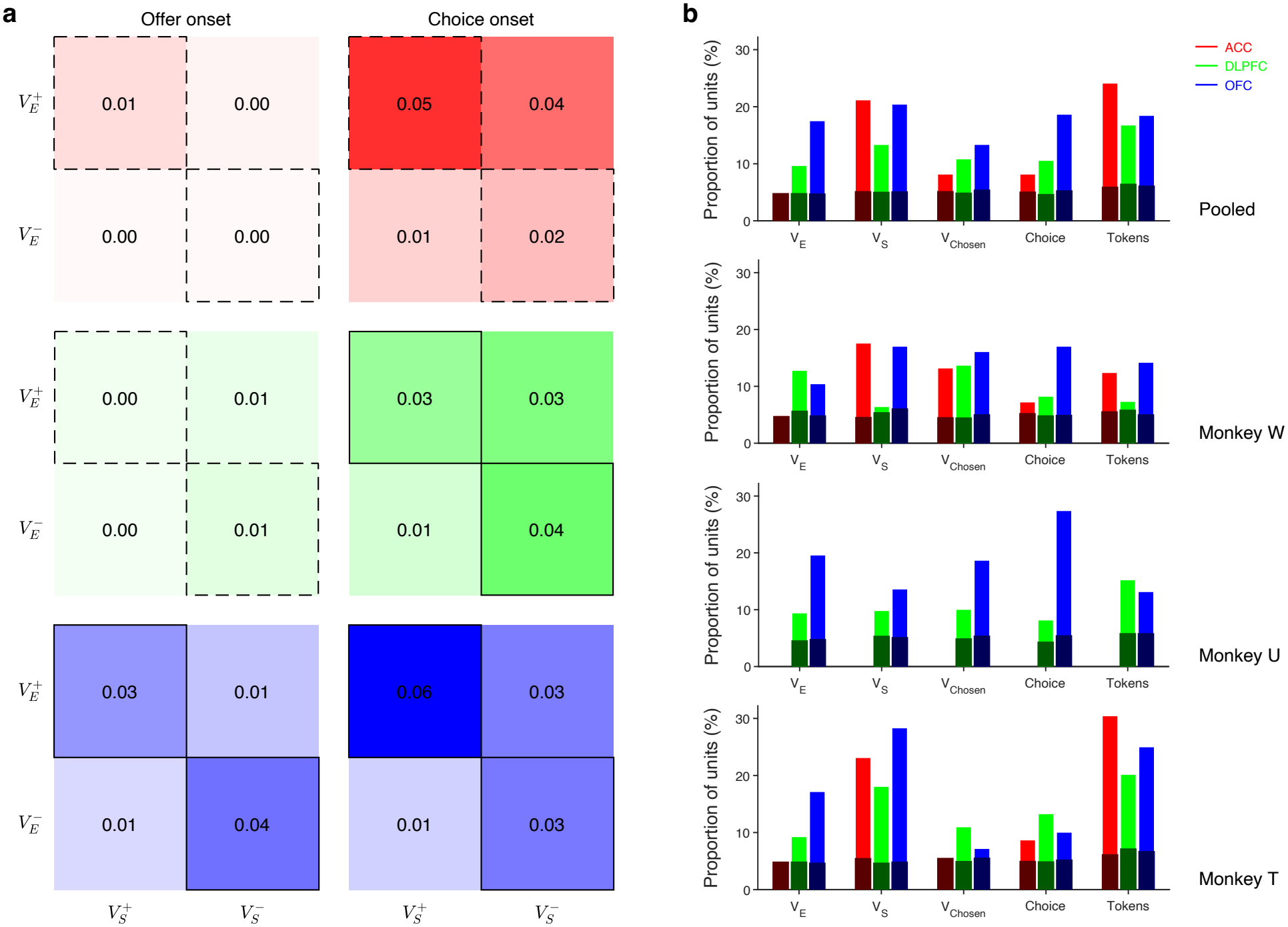
Single-unit decision-related value representations. **(a) Joint encoding of** *V*_*E*_ **and** *V*_*S*_. Coefficient-sign matrices show pooled results for ACC (red), DLPFC (green) and OFC (blue) in the offer-onset ([0, 500) ms; left) and choice-onset ([0, 150) ms; right) epochs. Rows indicate the sign of *V*_*E*_ tuning and columns the sign of *V*_*S*_ tuning. Values are proportions of all recorded units in each region. Borders mark diagonal associations (solid, significant; dashed, not significant). Two-sided Fisher’s exact tests yielded *p* = 1.00, 0.659 and 1.11 × 10^−4^ for ACC, DLPFC and OFC in the offer-onset epoch, and *p* = 0.198, 0.0192 and 1.47 × 10^−5^ in the choice-onset epoch, respectively. **(b) Proportions of variable-selective units**. Bars show the proportions of units encoding *V*_*E*_, *V*_*S*_, *V*_Chosen_, *Choice*_ES_ and token holdings in the offer-onset epoch ([0, 500) ms) in the pooled data and in monkeys W, U and T. Dark bars show the shuffle control; ACC was not recorded in monkey U.

## Supplementary Tables

**Supplementary Table 1.** Individual-animal and label-shuffle tests of choice-dependent updating of token-holding representations. The temporal coefficient of the token-holding-axis projection was tested against zero using two-sided Wilcoxon signed-rank tests; *p*_*E*_ and *p*_*S*_ denote the earning and spending tests, respectively. Earn *>* 0 and Spend *<* 0 give the percentages of resampling iterations with positive earning and negative spending coefficients, respectively. Significant *p* values (*p <* 0.05) are bold and colored by the sign of the median coefficient (red, positive; green, negative); non-significant values remain black. Significant coefficients were predominantly positive for earning and negative for spending. In DLPFC, only earning coefficients reached significance, possibly suggesting asymmetric updating. No ACC recordings were available for monkey U.

| Region | Dataset | Earn > 0 | *p*<sub>*E*</sub> | Spend < 0 | *p*<sub>*S*</sub> |
|---|---|---|---|---|---|
| ACC | Pooled data | 61% | **6.98e-04** | 74% | **9.55e-10** |
|  | Monkey W | 42% | **7.10e-03** | 47% | 7.67e-01 |
|  | Monkey T | 77% | **8.47e-08** | 72% | **8.49e-07** |
|  | Choice-label shuffle | 44% | 7.60e-02 | 58% | 1.31e-01 |
|  | Token-holding-label shuffle | 47% | 3.09e-01 | 52% | 8.96e-01 |
| DLPFC | Pooled data | 64% | **8.19e-03** | 59% | 1.12e-01 |
|  | Monkey W | 63% | **3.31e-04** | 44% | 6.87e-01 |
|  | Monkey U | 64% | **1.30e-02** | 45% | 8.07e-01 |
|  | Monkey T | 60% | **1.22e-02** | 56% | 1.45e-01 |
|  | Choice-label shuffle | 38% | 8.25e-02 | 48% | 9.62e-01 |
|  | Token-holding-label shuffle | 48% | 9.31e-01 | 54% | 5.80e-01 |
| OFC | Pooled data | 96% | **6.54e-17** | 69% | **1.81e-04** |
|  | Monkey W | 68% | **1.21e-03** | 66% | **4.36e-04** |
|  | Monkey U | 85% | **1.75e-14** | 75% | **4.32e-08** |
|  | Monkey T | 65% | **4.96e-04** | 53% | 2.51e-01 |
|  | Choice-label shuffle | 54% | 2.79e-01 | 40% | **2.24e-02** |
|  | Token-holding-label shuffle | 39% | 5.91e-02 | 55% | 1.74e-01 |

**Supplementary Table 2.**
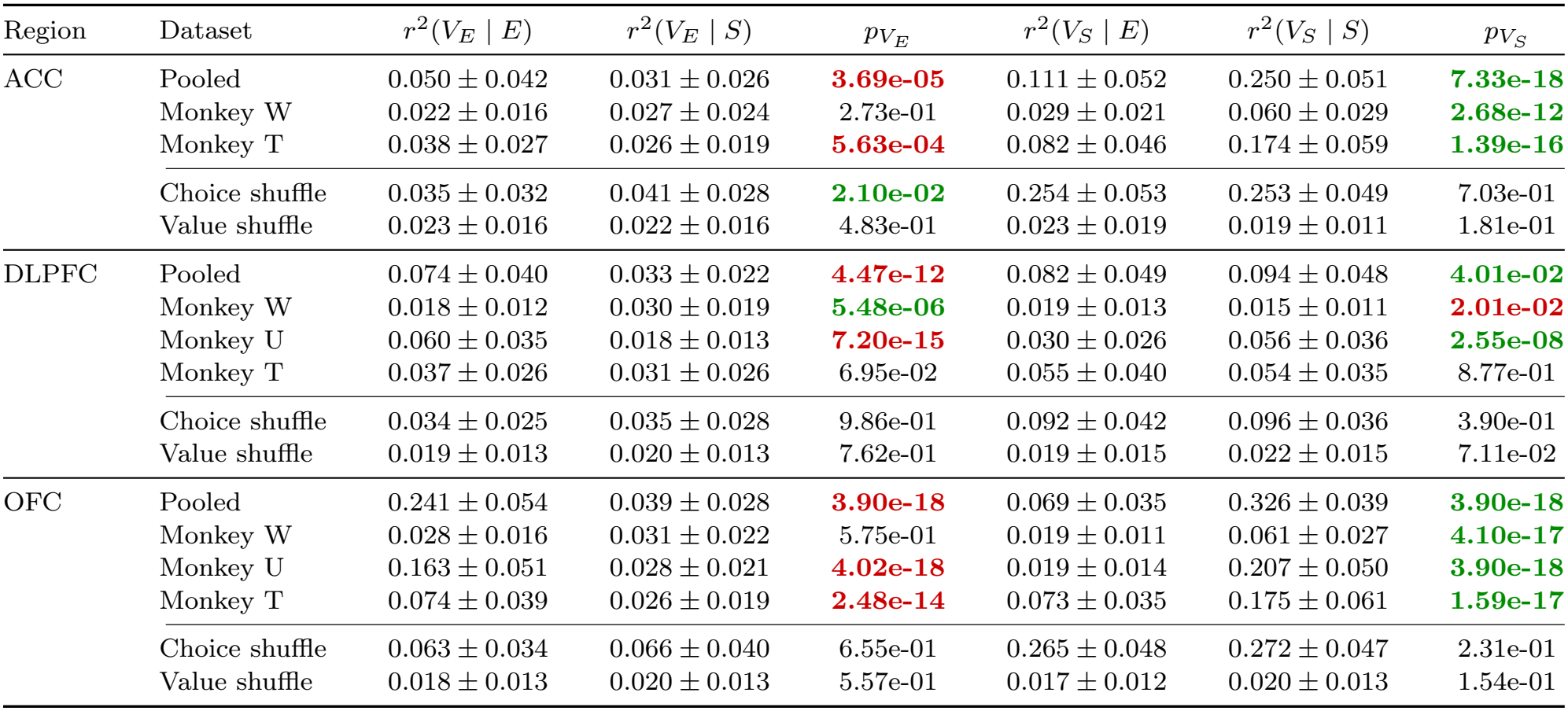
Choice-conditioned value encoding. Values are mean ± SD of squared Pearson correlations over [0, 500) ms after offer onset across 100 iterations. E and S denote earning and spending choices. Two-sided Wilcoxon signed-rank tests compared 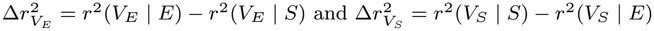 against zero. Significant *p* values are bold and colored by the direction of the difference: red indicates higher encoding strength on earning trials, and green indicates higher encoding strength on spending trials. Nonsignificant *p* values are black. Choice shuffle and Value shuffle denote the two label-shuffle controls in the pooled populations. Some significant differences opposite to the overall direction occurred at low absolute encoding strengths. These differences should be interpreted alongside the absolute encoding strengths and the magnitude of the differences, as small *p* values alone do not imply strong choice-dependent encoding. ACC was not recorded in monkey U.

| Region | Dataset | r<sup>2</sup>(V<sub>E</sub> \| E) | r<sup>2</sup>(V<sub>E</sub> \| S) | p<sub>V<sub>E</sub></sub> | r<sup>2</sup>(V<sub>S</sub> \| E) | r<sup>2</sup>(V<sub>S</sub> \| S) | p<sub>V<sub>S</sub></sub> |
|---|---|---|---|---|---|---|---|
| ACC | Pooled | 0.050 ± 0.042 | 0.031 ± 0.026 | **3.69e-05** | 0.111 ± 0.052 | 0.250 ± 0.051 | **7.33e-18** |
|  | Monkey W | 0.022 ± 0.016 | 0.027 ± 0.024 | 2.73e-01 | 0.029 ± 0.021 | 0.060 ± 0.029 | **2.68e-12** |
|  | Monkey T | 0.038 ± 0.027 | 0.026 ± 0.019 | **5.63e-04** | 0.082 ± 0.046 | 0.174 ± 0.059 | **1.39e-16** |
|  | Choice shuffle | 0.035 ± 0.032 | 0.041 ± 0.028 | **2.10e-02** | 0.254 ± 0.053 | 0.253 ± 0.049 | 7.03e-01 |
|  | Value shuffle | 0.023 ± 0.016 | 0.022 ± 0.016 | 4.83e-01 | 0.023 ± 0.019 | 0.019 ± 0.011 | 1.81e-01 |
| DLPFC | Pooled | 0.074 ± 0.040 | 0.033 ± 0.022 | **4.47e-12** | 0.082 ± 0.049 | 0.094 ± 0.048 | **4.01e-02** |
|  | Monkey W | 0.018 ± 0.012 | 0.030 ± 0.019 | **5.48e-06** | 0.019 ± 0.013 | 0.015 ± 0.011 | **2.01e-02** |
|  | Monkey U | 0.060 ± 0.035 | 0.018 ± 0.013 | **7.20e-15** | 0.030 ± 0.026 | 0.056 ± 0.036 | **2.55e-08** |
|  | Monkey T | 0.037 ± 0.026 | 0.031 ± 0.026 | 6.95e-02 | 0.055 ± 0.040 | 0.054 ± 0.035 | 8.77e-01 |
|  | Choice shuffle | 0.034 ± 0.025 | 0.035 ± 0.028 | 9.86e-01 | 0.092 ± 0.042 | 0.096 ± 0.036 | 3.90e-01 |
|  | Value shuffle | 0.019 ± 0.013 | 0.020 ± 0.013 | 7.62e-01 | 0.019 ± 0.015 | 0.022 ± 0.015 | 7.11e-02 |
| OFC | Pooled | 0.241 ± 0.054 | 0.039 ± 0.028 | **3.90e-18** | 0.069 ± 0.035 | 0.326 ± 0.039 | **3.90e-18** |
|  | Monkey W | 0.028 ± 0.016 | 0.031 ± 0.022 | 5.75e-01 | 0.019 ± 0.011 | 0.061 ± 0.027 | **4.10e-17** |
|  | Monkey U | 0.163 ± 0.051 | 0.028 ± 0.021 | **4.02e-18** | 0.019 ± 0.014 | 0.207 ± 0.050 | **3.90e-18** |
|  | Monkey T | 0.074 ± 0.039 | 0.026 ± 0.019 | **2.48e-14** | 0.073 ± 0.035 | 0.175 ± 0.061 | **1.59e-17** |
|  | Choice shuffle | 0.063 ± 0.034 | 0.066 ± 0.040 | 6.55e-01 | 0.265 ± 0.048 | 0.272 ± 0.047 | 2.31e-01 |
|  | Value shuffle | 0.018 ± 0.013 | 0.020 ± 0.013 | 5.57e-01 | 0.017 ± 0.012 | 0.020 ± 0.013 | 1.54e-01 |

**Supplementary Table 3.**
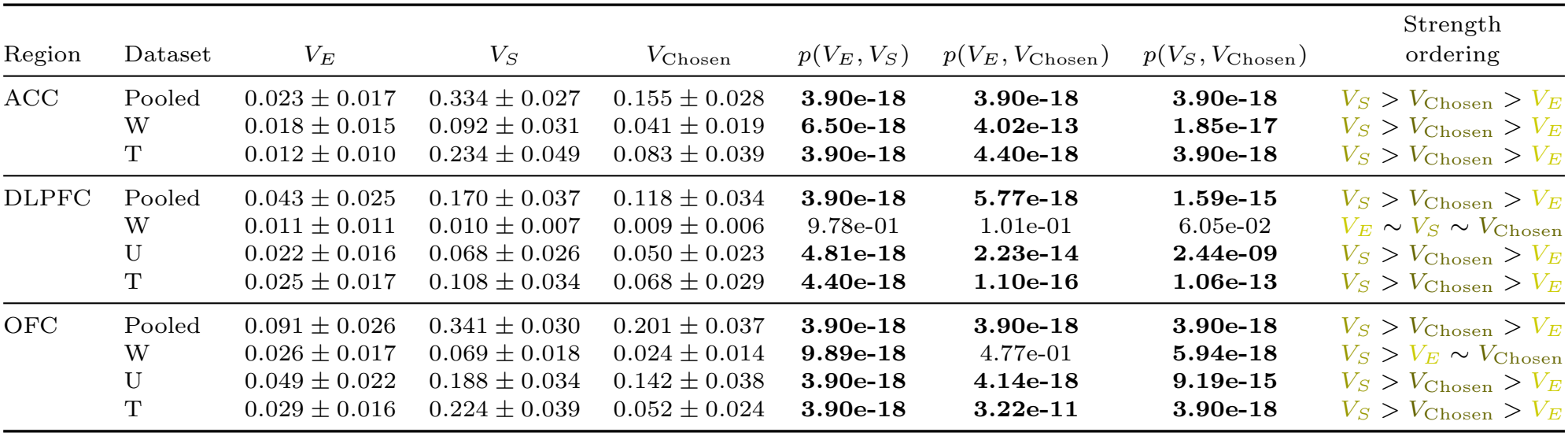
Encoding-strength comparisons within brain area. Mean ± SD of *r*^2^ over 0–500 ms across 100 iterations. Two-sided paired Wilcoxon signed-rank tests; significant *p* values are bold. Strength ordering ranks the means; ∼ denotes a nonsignificant adjacent comparison, not equivalence. Ordering colors identify *V*_*E*_, *V*_*S*_ and *V*_Chosen_. ACC was not recorded in monkey U.

**Supplementary Table 4.**
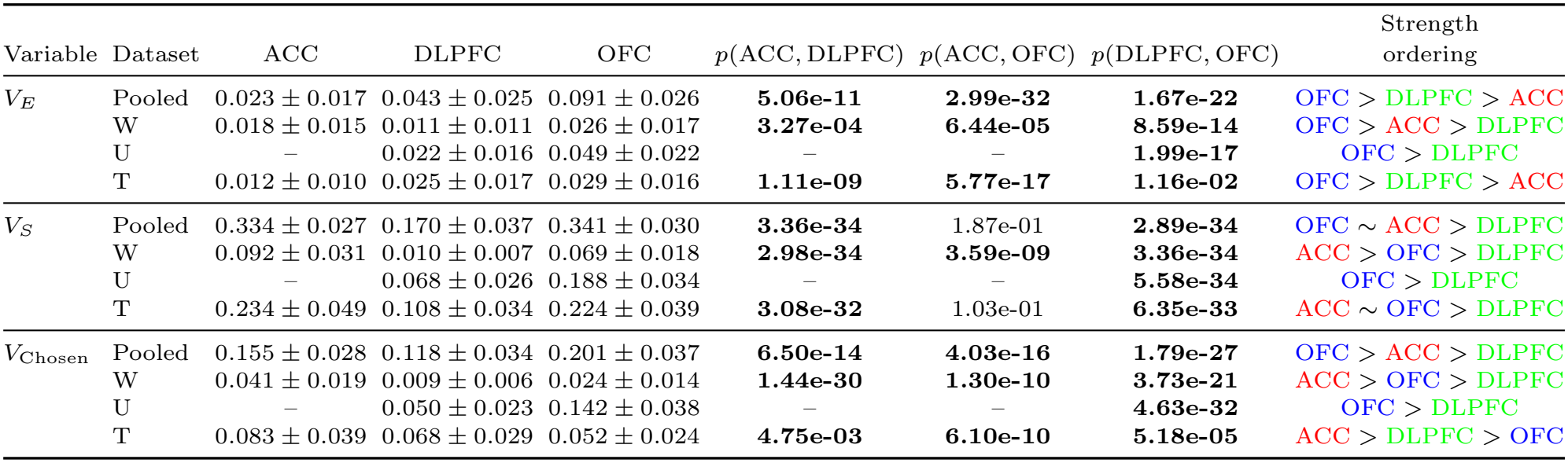
Encoding-strength comparisons between brain areas. Mean ± SD of *r*^2^ over 0–500 ms across 100 iterations. Two-sided Wilcoxon rank-sum tests; significant *p* values are bold. Strength ordering ranks the means; ∼ denotes a nonsignificant adjacent comparison, not equivalence. Ordering colors identify ACC, DLPFC and OFC in red, green and blue, respectively. ACC was not recorded in monkey U.

| Variable | Dataset | ACC | DLPFC | OFC | *p*(ACC, DLPFC) | *p*(ACC, OFC) | *p*(DLPFC, OFC) | Strength ordering |
|---|---|---|---|---|---|---|---|---|
| *V<sub>E</sub>* | Pooled | 0.023 ± 0.017 | 0.043 ± 0.025 | 0.091 ± 0.026 | **5.06e-11** | **2.99e-32** | **1.67e-22** | OFC > DLPFC > ACC |
|  | W | 0.018 ± 0.015 | 0.011 ± 0.011 | 0.026 ± 0.017 | **3.27e-04** | **6.44e-05** | **8.59e-14** | OFC > ACC > DLPFC |
|  | U | – | 0.022 ± 0.016 | 0.049 ± 0.022 | – | – | **1.99e-17** | OFC > DLPFC |
|  | T | 0.012 ± 0.010 | 0.025 ± 0.017 | 0.029 ± 0.016 | **1.11e-09** | **5.77e-17** | **1.16e-02** | OFC > DLPFC > ACC |
| *V<sub>S</sub>* | Pooled | 0.334 ± 0.027 | 0.170 ± 0.037 | 0.341 ± 0.030 | **3.36e-34** | 1.87e-01 | **2.89e-34** | OFC ∼ ACC > DLPFC |
|  | W | 0.092 ± 0.031 | 0.010 ± 0.007 | 0.069 ± 0.018 | **2.98e-34** | **3.59e-09** | **3.36e-34** | ACC > OFC > DLPFC |
|  | U | – | 0.068 ± 0.026 | 0.188 ± 0.034 | – | – | **5.58e-34** | OFC > DLPFC |
|  | T | 0.234 ± 0.049 | 0.108 ± 0.034 | 0.224 ± 0.039 | **3.08e-32** | 1.03e-01 | **6.35e-33** | ACC ∼ OFC > DLPFC |
| *V*<sub>Chosen</sub> | Pooled | 0.155 ± 0.028 | 0.118 ± 0.034 | 0.201 ± 0.037 | **6.50e-14** | **4.03e-16** | **1.79e-27** | OFC > ACC > DLPFC |
|  | W | 0.041 ± 0.019 | 0.009 ± 0.006 | 0.024 ± 0.014 | **1.44e-30** | **1.30e-10** | **3.73e-21** | ACC > OFC > DLPFC |
|  | U | – | 0.050 ± 0.023 | 0.142 ± 0.038 | – | – | **4.63e-32** | OFC > DLPFC |
|  | T | 0.083 ± 0.039 | 0.068 ± 0.029 | 0.052 ± 0.024 | **4.75e-03** | **6.10e-10** | **5.18e-05** | ACC > DLPFC > OFC |

**Supplementary Table 5.** Half-peak latency (*t*_50_) comparisons within brain area. Mean ± SD (ms) and valid percentage across 100 iterations. Valid (%) is the percentage of the 100 iterations that yielded a latency estimate (see Methods, Latency). Two-sided Wilcoxon rank-sum tests compare each signal with *Choice*_ES_ and *V*_Chosen_; bold red/green indicates an earlier/later signal. Dashes indicate inapplicable comparisons. A minority of comparisons showed significant differences opposite to the overall temporal order. These comparisons should be interpreted in light of the corresponding encoding strengths, valid percentages and latency SDs. When encoding remains close to the pre-offer baseline, latency estimates are unstable, and even a small *p* value does not establish a reliable temporal ordering. ACC was not recorded in monkey U.

| Region | Dataset | Variable | Mean ± SD (ms) | Valid (%) | *p* vs *Choice*<sub>ES</sub> | *p* vs *V*<sub>Chosen</sub> |
|---|---|---|---|---|---|---|
| ACC | Pooled | *V*<sub>*E*</sub> | 194.5 ± 130.4 | 87 | **4.74e-09** | **5.52e-07** |
|  |  | *V*<sub>*S*</sub> | 267.1 ± 10.3 | 100 | **3.44e-11** | **2.18e-14** |
|  |  | *V*<sub>Chosen</sub> | 287.8 ± 22.0 | 100 | **1.74e-03** | – |
|  |  | *Choice*<sub>ES</sub> | 302.7 ± 68.2 | 99 | – | – |
|  | Monkey W | *V*<sub>*E*</sub> | 153.5 ± 133.4 | 81 | **1.58e-13** | **6.01e-15** |
|  |  | *V*<sub>*S*</sub> | 327.1 ± 19.4 | 100 | 1.61e-01 | 5.53e-02 |
|  |  | *V*<sub>Chosen</sub> | 324.3 ± 82.7 | 98 | 9.11e-01 | – |
|  |  | *Choice*<sub>ES</sub> | 320.0 ± 102.5 | 94 | – | – |
|  | Monkey T | *V*<sub>*E*</sub> | 177.0 ± 132.3 | 83 | **1.07e-05** | **1.04e-09** |
|  |  | *V*<sub>*S*</sub> | 276.5 ± 16.8 | 100 | 2.76e-01 | **9.96e-03** |
|  |  | *V*<sub>Chosen</sub> | 289.0 ± 51.7 | 98 | 5.85e-01 | – |
|  |  | *Choice*<sub>ES</sub> | 263.9 ± 110.5 | 90 | – | – |
| DLPFC | Pooled | *V*<sub>*E*</sub> | 296.3 ± 88.6 | 98 | **4.56e-09** | 1.94e-01 |
|  |  | *V*<sub>*S*</sub> | 298.6 ± 16.8 | 100 | **1.70e-23** | **8.23e-05** |
|  |  | *V*<sub>Chosen</sub> | 311.3 ± 26.3 | 100 | **1.68e-14** | – |
|  |  | *Choice*<sub>ES</sub> | 344.6 ± 44.3 | 100 | – | – |
|  | Monkey W | *V*<sub>*E*</sub> | 226.0 ± 154.6 | 84 | 6.15e-01 | **2.42e-02** |
|  |  | *V*<sub>*S*</sub> | 187.6 ± 146.1 | 86 | 5.99e-01 | **9.62e-04** |
|  |  | *V*<sub>Chosen</sub> | 278.9 ± 169.0 | 74 | **1.66e-02** | – |
|  |  | *Choice*<sub>ES</sub> | 217.7 ± 168.3 | 59 | – | – |
|  | Monkey U | *V*<sub>*E*</sub> | 252.4 ± 103.7 | 92 | **3.05e-12** | **5.54e-11** |
|  |  | *V*<sub>*S*</sub> | 331.0 ± 46.4 | 100 | **3.80e-03** | 3.26e-01 |
|  |  | *V*<sub>Chosen</sub> | 325.5 ± 56.3 | 100 | **1.23e-03** | – |
|  |  | *Choice*<sub>ES</sub> | 339.3 ± 104.5 | 94 | – | – |
|  | Monkey T | *V*<sub>*E*</sub> | 260.0 ± 142.3 | 94 | **3.09e-04** | 3.87e-01 |
|  |  | *V*<sub>*S*</sub> | 293.1 ± 28.9 | 100 | **1.28e-16** | **5.20e-04** |
|  |  | *V*<sub>Chosen</sub> | 304.1 ± 66.4 | 100 | **6.85e-08** | – |
|  |  | *Choice*<sub>ES</sub> | 333.9 ± 93.0 | 99 | – | – |
| OFC | Pooled | *V*<sub>*E*</sub> | 259.5 ± 53.2 | 100 | **1.10e-13** | 7.73e-01 |
|  |  | *V*<sub>*S*</sub> | 239.8 ± 11.3 | 100 | **1.96e-33** | **1.86e-14** |
|  |  | *V*<sub>Chosen</sub> | 261.8 ± 22.1 | 100 | **4.76e-24** | – |
|  |  | *Choice*<sub>ES</sub> | 304.6 ± 21.5 | 100 | – | – |
|  | Monkey W | *V*<sub>*E*</sub> | 236.5 ± 133.0 | 84 | **9.40e-09** | **7.09e-04** |
|  |  | *V*<sub>*S*</sub> | 374.6 ± 24.7 | 100 | **1.80e-03** | **4.22e-09** |
|  |  | *V*<sub>Chosen</sub> | 297.0 ± 112.6 | 98 | **8.01e-03** | – |
|  |  | *Choice*<sub>ES</sub> | 342.8 ± 83.4 | 100 | – | – |
|  | Monkey U | *V*<sub>*E*</sub> | 262.6 ± 57.6 | 99 | **8.37e-15** | 7.03e-02 |
|  |  | *V*<sub>*S*</sub> | 264.4 ± 17.5 | 100 | **2.69e-30** | **2.70e-03** |
|  |  | *V*<sub>Chosen</sub> | 272.2 ± 19.4 | 100 | **2.81e-26** | – |
|  |  | *Choice*<sub>ES</sub> | 314.8 ± 21.9 | 100 | – | – |
|  | Monkey T | *V*<sub>*E*</sub> | 255.8 ± 117.6 | 89 | 4.24e-01 | 8.23e-01 |
|  |  | *V*<sub>*S*</sub> | 242.8 ± 15.0 | 100 | **4.30e-04** | **1.07e-06** |
|  |  | *V*<sub>Chosen</sub> | 273.0 ± 81.3 | 96 | 5.15e-01 | – |
|  |  | *Choice*<sub>ES</sub> | 269.0 ± 126.8 | 87 | – | – |

**Supplementary Table 6.** Onset-latency comparisons within brain area. Mean ± SD (ms) and valid percentage across 100 iterations. Valid (%) is the percentage of the 100 iterations that yielded a latency estimate (see Methods, Latency). Two-sided Wilcoxon rank-sum tests compare each signal with *Choice*_ES_ and *V*_Chosen_; bold red/green indicates an earlier/later signal. Dashes indicate inapplicable comparisons. A minority of comparisons showed significant differences opposite to the overall temporal order. These comparisons should be interpreted in light of the corresponding encoding strengths, valid percentages and latency SDs. When encoding remains close to the pre-offer baseline, latency estimates are unstable, and even a small *p* value does not establish a reliable temporal ordering. ACC was not recorded in monkey U.

| Region | Dataset | Variable | Mean ± SD (ms) | Valid (%) | *p* vs *Choice*<sub>ES</sub> | *p* vs *V*<sub>Chosen</sub> |
|---|---|---|---|---|---|---|
| ACC | Pooled | *V*<sub>*E*</sub> | 173.9 ± 121.1 | 73 | **2.11e-02** | 2.68e-01 |
|  |  | *V*<sub>*S*</sub> | 145.3 ± 69.4 | 100 | **2.21e-08** | 6.00e-01 |
|  |  | *V*<sub>Chosen</sub> | 141.6 ± 82.9 | 100 | **6.82e-08** | – |
|  |  | *Choice*<sub>ES</sub> | 210.8 ± 101.6 | 99 | – | – |
|  | Monkey W | *V*<sub>*E*</sub> | 133.5 ± 122.0 | 55 | **4.24e-06** | **7.02e-06** |
|  |  | *V*<sub>*S*</sub> | 203.9 ± 102.1 | 99 | **2.27e-02** | 7.09e-02 |
|  |  | *V*<sub>Chosen</sub> | 224.6 ± 107.1 | 96 | 4.94e-01 | – |
|  |  | *Choice*<sub>ES</sub> | 235.3 ± 117.4 | 91 | – | – |
|  | Monkey T | *V*<sub>*E*</sub> | 181.6 ± 123.6 | 59 | 9.32e-02 | 3.50e-01 |
|  |  | *V*<sub>*S*</sub> | 141.5 ± 78.7 | 100 | **3.96e-05** | **2.02e-02** |
|  |  | *V*<sub>Chosen</sub> | 163.1 ± 95.5 | 97 | **2.62e-03** | – |
|  |  | *Choice*<sub>ES</sub> | 215.0 ± 116.5 | 82 | – | – |
| DLPFC | Pooled | *V*<sub>*E*</sub> | 209.2 ± 117.6 | 97 | 7.69e-02 | **1.09e-03** |
|  |  | *V*<sub>*S*</sub> | 175.9 ± 78.2 | 100 | **3.35e-11** | 2.69e-01 |
|  |  | *V*<sub>Chosen</sub> | 170.1 ± 78.8 | 100 | **5.61e-10** | – |
|  |  | *Choice*<sub>ES</sub> | 233.0 ± 88.1 | 99 | – | – |
|  | Monkey W | *V*<sub>*E*</sub> | 187.0 ± 134.8 | 61 | 5.86e-01 | **4.21e-03** |
|  |  | *V*<sub>*S*</sub> | 185.6 ± 135.1 | 62 | 6.91e-01 | **4.39e-03** |
|  |  | *V*<sub>Chosen</sub> | 262.5 ± 145.9 | 52 | 5.52e-02 | – |
|  |  | *Choice*<sub>ES</sub> | 207.9 ± 146.3 | 38 | – | – |
|  | Monkey U | *V*<sub>*E*</sub> | 204.7 ± 106.4 | 78 | 3.37e-01 | 6.49e-01 |
|  |  | *V*<sub>*S*</sub> | 199.4 ± 102.3 | 100 | 1.54e-01 | 4.04e-01 |
|  |  | *V*<sub>Chosen</sub> | 213.2 ± 93.5 | 99 | 5.13e-01 | – |
|  |  | *Choice*<sub>ES</sub> | 216.5 ± 119.2 | 92 | – | – |
|  | Monkey T | *V*<sub>*E*</sub> | 228.6 ± 133.1 | 81 | 7.81e-01 | **2.98e-02** |
|  |  | *V*<sub>*S*</sub> | 179.4 ± 89.7 | 100 | **4.01e-05** | 2.04e-01 |
|  |  | *V*<sub>Chosen</sub> | 195.9 ± 97.8 | 100 | **1.29e-02** | – |
|  |  | *Choice*<sub>ES</sub> | 223.8 ± 125.4 | 96 | – | – |
| OFC | Pooled | *V*<sub>*E*</sub> | 176.4 ± 71.2 | 100 | 7.95e-02 | 5.21e-01 |
|  |  | *V*<sub>*S*</sub> | 162.4 ± 54.2 | 100 | **3.84e-09** | **4.62e-04** |
|  |  | *V*<sub>Chosen</sub> | 174.3 ± 54.6 | 100 | **2.06e-04** | – |
|  |  | *Choice*<sub>ES</sub> | 190.4 ± 62.1 | 100 | – | – |
|  | Monkey W | *V*<sub>*E*</sub> | 229.6 ± 121.5 | 69 | 9.06e-01 | 4.31e-01 |
|  |  | *V*<sub>*S*</sub> | 245.9 ± 103.0 | 100 | 3.46e-01 | **3.76e-02** |
|  |  | *V*<sub>Chosen</sub> | 216.3 ± 117.8 | 94 | 4.38e-01 | – |
|  |  | *Choice*<sub>ES</sub> | 227.4 ± 133.0 | 99 | – | – |
|  | Monkey U | *V*<sub>*E*</sub> | 170.2 ± 83.7 | 99 | 5.45e-01 | 3.95e-01 |
|  |  | *V*<sub>*S*</sub> | 167.1 ± 62.8 | 100 | **4.85e-02** | **3.54e-02** |
|  |  | *V*<sub>Chosen</sub> | 174.9 ± 63.7 | 100 | 5.67e-01 | – |
|  |  | *Choice*<sub>ES</sub> | 169.8 ± 75.8 | 100 | – | – |
|  | Monkey T | *V*<sub>*E*</sub> | 204.7 ± 119.1 | 79 | 2.30e-01 | 7.53e-01 |
|  |  | *V*<sub>*S*</sub> | 159.0 ± 67.8 | 100 | **6.46e-06** | **4.14e-07** |
|  |  | *V*<sub>Chosen</sub> | 204.4 ± 93.9 | 86 | 1.48e-01 | – |
|  |  | *Choice*<sub>ES</sub> | 229.1 ± 129.4 | 73 | – | – |

**Supplementary Table 7.**
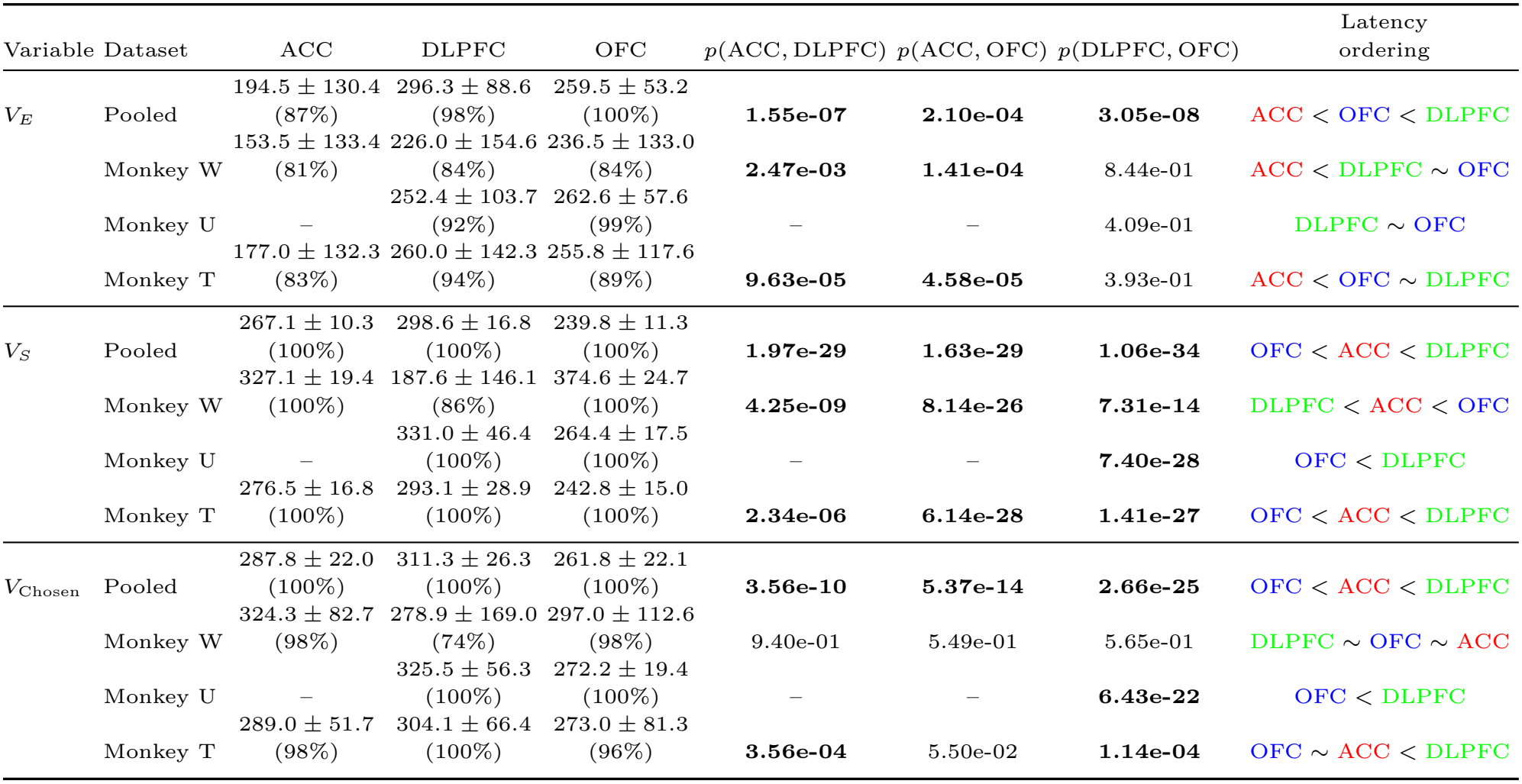
Half-peak latency (*t*_50_) comparisons between brain areas. Mean ± SD (ms), with valid percentage in parentheses. Valid percentage is the percentage of the 100 iterations that yielded a latency estimate (see Methods, Latency). Two-sided Wilcoxon rank-sum tests; significant *p* values are bold. Latency ordering ranks the means; *<* denotes an earlier signal and ∼ a nonsignificant adjacent comparison, not equivalence. ACC was not recorded in monkey U.

| Variable | Dataset | ACC | DLPFC | OFC | *p*(ACC, DLPFC) | *p*(ACC, OFC) | *p*(DLPFC, OFC) | Latency<br>ordering |
|---|---|---|---|---|---|---|---|---|
| *V*<sub>*E*</sub> | Pooled | 194.5 ± 130.4<br>(87%) | 296.3 ± 88.6<br>(98%) | 259.5 ± 53.2<br>(100%) | **1.55e-07** | **2.10e-04** | **3.05e-08** | ACC < OFC < DLPFC |
|  | Monkey W | 153.5 ± 133.4<br>(81%) | 226.0 ± 154.6<br>(84%) | 236.5 ± 133.0<br>(84%) | **2.47e-03** | **1.41e-04** | 8.44e-01 | ACC < DLPFC ∼ OFC |
|  | Monkey U | – | 252.4 ± 103.7<br>(92%) | 262.6 ± 57.6<br>(99%) | – | – | 4.09e-01 | DLPFC ∼ OFC |
|  | Monkey T | 177.0 ± 132.3<br>(83%) | 260.0 ± 142.3<br>(94%) | 255.8 ± 117.6<br>(89%) | **9.63e-05** | **4.58e-05** | 3.93e-01 | ACC < OFC ∼ DLPFC |
| *V*<sub>*S*</sub> | Pooled | 267.1 ± 10.3<br>(100%) | 298.6 ± 16.8<br>(100%) | 239.8 ± 11.3<br>(100%) | **1.97e-29** | **1.63e-29** | **1.06e-34** | OFC < ACC < DLPFC |
|  | Monkey W | 327.1 ± 19.4<br>(100%) | 187.6 ± 146.1<br>(86%) | 374.6 ± 24.7<br>(100%) | **4.25e-09** | **8.14e-26** | **7.31e-14** | DLPFC < ACC < OFC |
|  | Monkey U | – | 331.0 ± 46.4<br>(100%) | 264.4 ± 17.5<br>(100%) | – | – | **7.40e-28** | OFC < DLPFC |
|  | Monkey T | 276.5 ± 16.8<br>(100%) | 293.1 ± 28.9<br>(100%) | 242.8 ± 15.0<br>(100%) | **2.34e-06** | **6.14e-28** | **1.41e-27** | OFC < ACC < DLPFC |
| *V*<sub>Chosen</sub> | Pooled | 287.8 ± 22.0<br>(100%) | 311.3 ± 26.3<br>(100%) | 261.8 ± 22.1<br>(100%) | **3.56e-10** | **5.37e-14** | **2.66e-25** | OFC < ACC < DLPFC |
|  | Monkey W | 324.3 ± 82.7<br>(98%) | 278.9 ± 169.0<br>(74%) | 297.0 ± 112.6<br>(98%) | 9.40e-01 | 5.49e-01 | 5.65e-01 | DLPFC ∼ OFC ∼ ACC |
|  | Monkey U | – | 325.5 ± 56.3<br>(100%) | 272.2 ± 19.4<br>(100%) | – | – | **6.43e-22** | OFC < DLPFC |
|  | Monkey T | 289.0 ± 51.7<br>(98%) | 304.1 ± 66.4<br>(100%) | 273.0 ± 81.3<br>(96%) | **3.56e-04** | 5.50e-02 | **1.14e-04** | OFC ∼ ACC < DLPFC |

**Supplementary Table 8.** Onset-latency comparisons between brain areas. Mean ± SD (ms), with valid percentage in parentheses. Valid percentage is the percentage of the 100 iterations that yielded a latency estimate (see Methods, Latency). Two-sided Wilcoxon rank-sum tests; significant *p* values are bold. Latency ordering ranks the means; *<* denotes an earlier signal and ∼ a nonsignificant adjacent comparison, not equivalence. ACC was not recorded in monkey U.

| Variable | Dataset | ACC | DLPFC | OFC | *p*(ACC, DLPFC) | *p*(ACC, OFC) | *p*(DLPFC, OFC) | Latency<br>ordering |
|---|---|---|---|---|---|---|---|---|
| *V<sub>E</sub>* | Pooled | 173.9 ± 121.1<br>(73%) | 209.2 ± 117.6<br>(97%) | 176.4 ± 71.2<br>(100%) | **4.95e-02** | 2.47e-01 | **1.35e-03** | ACC ∼ OFC < DLPFC |
|  | Monkey W | 133.5 ± 122.0<br>(55%) | 187.0 ± 134.8<br>(61%) | 229.6 ± 121.5<br>(69%) | **3.94e-02** | **3.88e-05** | 8.33e-02 | ACC < DLPFC ∼ OFC |
|  | Monkey U | – | 204.7 ± 106.4<br>(78%) | 170.2 ± 83.7<br>(99%) | – | – | **2.12e-03** | OFC < DLPFC |
|  | Monkey T | 181.6 ± 123.6<br>(59%) | 228.6 ± 133.1<br>(81%) | 204.7 ± 119.1<br>(79%) | **3.66e-02** | 2.47e-01 | 1.93e-01 | ACC ∼ OFC ∼ DLPFC |
| *V<sub>S</sub>* | Pooled | 145.3 ± 69.4<br>(100%) | 175.9 ± 78.2<br>(100%) | 162.4 ± 54.2<br>(100%) | **6.51e-06** | 8.72e-01 | **1.94e-06** | ACC ∼ OFC < DLPFC |
|  | Monkey W | 203.9 ± 102.1<br>(99%) | 185.6 ± 135.1<br>(62%) | 245.9 ± 103.0<br>(100%) | 2.13e-01 | **3.27e-05** | **4.83e-03** | DLPFC ∼ ACC < OFC |
|  | Monkey U | – | 199.4 ± 102.3<br>(100%) | 167.1 ± 62.8<br>(100%) | – | – | **5.85e-05** | OFC < DLPFC |
|  | Monkey T | 141.5 ± 78.7<br>(100%) | 179.4 ± 89.7<br>(100%) | 159.0 ± 67.8<br>(100%) | **9.45e-06** | 7.40e-01 | **1.12e-05** | ACC ∼ OFC < DLPFC |
| *V*<sub>Chosen</sub> | Pooled | 141.6 ± 82.9<br>(100%) | 170.1 ± 78.8<br>(100%) | 174.3 ± 54.6<br>(100%) | **2.84e-02** | 1.02e-01 | 7.70e-01 | ACC < DLPFC ∼ OFC |
|  | Monkey W | 224.6 ± 107.1<br>(96%) | 262.5 ± 145.9<br>(52%) | 216.3 ± 117.8<br>(94%) | 8.68e-02 | 4.73e-01 | **3.33e-02** | OFC ∼ ACC ∼ DLPFC |
|  | Monkey U | – | 213.2 ± 93.5<br>(99%) | 174.9 ± 63.7<br>(100%) | – | – | **9.26e-06** | OFC < DLPFC |
|  | Monkey T | 163.1 ± 95.5<br>(97%) | 195.9 ± 97.8<br>(100%) | 204.4 ± 93.9<br>(86%) | **2.31e-02** | **1.05e-02** | 6.19e-01 | ACC < DLPFC ∼ OFC |

**Supplementary Table 9.** Encoding-strength comparisons within brain area for good-based and action-based choice signals. Mean ± SD of *r*^2^ over [0, 500) ms after offer onset across 100 iterations for the earning/spending (ES) and left/right (LR) choice signals. Two-sided paired Wilcoxon signed-rank tests compare ES and LR signals; significant *p* values are bold. Strength ordering ranks the means; ∼ denotes a nonsignificant comparison, not equivalence. Ordering colors identify ES and LR in olive and blue, respectively. ACC was not recorded in monkey U.

| Region | Dataset | ES *r*<sup>2</sup> | LR *r*<sup>2</sup> | *p*(ES, LR) | Strength<br>ordering |
|---|---|---|---|---|---|
| ACC | Pooled | 0.056 ± 0.026 | 0.016 ± 0.012 | **3.98e-17** | ES > LR |
|  | Monkey W | 0.030 ± 0.018 | 0.010 ± 0.007 | **1.04e-15** | ES > LR |
|  | Monkey T | 0.031 ± 0.022 | 0.012 ± 0.010 | **1.13e-12** | ES > LR |
| DLPFC | Pooled | 0.092 ± 0.033 | 0.318 ± 0.033 | **3.90e-18** | LR > ES |
|  | Monkey W | 0.011 ± 0.007 | 0.008 ± 0.006 | **4.83e-04** | ES > LR |
|  | Monkey U | 0.041 ± 0.023 | 0.053 ± 0.022 | **1.45e-04** | LR > ES |
|  | Monkey T | 0.053 ± 0.027 | 0.288 ± 0.043 | **3.90e-18** | LR > ES |
| OFC | Pooled | 0.212 ± 0.037 | 0.084 ± 0.029 | **3.90e-18** | ES > LR |
|  | Monkey W | 0.030 ± 0.014 | 0.012 ± 0.009 | **3.44e-15** | ES > LR |
|  | Monkey U | 0.165 ± 0.034 | 0.067 ± 0.032 | **5.94e-18** | ES > LR |
|  | Monkey T | 0.038 ± 0.023 | 0.031 ± 0.024 | **7.95e-03** | ES > LR |

**Supplementary Table 10.** Onset-latency comparisons within brain area for good-based and action-based choice signals. Mean ± SD (ms), with valid percentage in parentheses, across 100 iterations for the earning/spending (ES) and left/right (LR) choice signals. Valid percentage is the percentage of the 100 iterations that yielded a latency estimate (see Methods, Latency). Two-sided Wilcoxon rank-sum tests compare ES and LR signals; significant *p* values are bold. Latency ordering ranks the means; *<* denotes an earlier signal and ∼ a nonsignificant comparison, not equivalence. Weak signals can yield unstable estimates. ACC was not recorded in monkey U.

| Region | Dataset | ES latency | LR latency | *p*(ES, LR) | Latency<br>ordering |
|---|---|---|---|---|---|
| ACC | Pooled | 216.8 ± 96.8<br>(100%) | 178.2 ± 135.6<br>(63%) | 6.26e-02 | LR ∼ ES |
|  | Monkey W | 240.7 ± 115.5<br>(93%) | 168.9 ± 139.1<br>(41%) | **6.76e-03** | LR < ES |
|  | Monkey T | 199.4 ± 122.7<br>(77%) | 226.9 ± 131.1<br>(64%) | 1.77e-01 | ES ∼ LR |
| DLPFC | Pooled | 215.9 ± 101.7<br>(100%) | 143.0 ± 75.0<br>(100%) | **1.57e-09** | LR < ES |
|  | Monkey W | 273.3 ± 119.3<br>(35%) | 184.8 ± 148.1<br>(50%) | **5.16e-03** | LR < ES |
|  | Monkey U | 242.6 ± 104.3<br>(96%) | 223.8 ± 115.0<br>(100%) | 2.77e-01 | LR ∼ ES |
|  | Monkey T | 212.9 ± 128.0<br>(94%) | 143.6 ± 72.3<br>(100%) | **3.73e-06** | LR < ES |
| OFC | Pooled | 195.5 ± 63.2<br>(100%) | 176.6 ± 72.3<br>(100%) | **1.66e-02** | LR < ES |
|  | Monkey W | 246.3 ± 123.2<br>(92%) | 195.1 ± 145.6<br>(69%) | **1.34e-02** | LR < ES |
|  | Monkey U | 159.5 ± 84.6<br>(100%) | 187.0 ± 64.1<br>(98%) | **4.68e-02** | ES < LR |
|  | Monkey T | 222.3 ± 121.2<br>(67%) | 218.1 ± 112.1<br>(81%) | 7.52e-01 | LR ∼ ES |

**Supplementary Table 11.** Half-peak latency (*t*_50_) comparisons within brain area for good-based and action-based choice signals. ES and LR denote earning/spending and left/right choice signals, respectively. Same conventions as Supplementary Table 10, except that latency is quantified by *t*_50_.

| Region | Dataset | ES *t*<sub>50</sub> | LR *t*<sub>50</sub> | *p*(ES, LR) | Latency ordering |
|---|---|---|---|---|---|
| ACC | Pooled | 311.2 ± 72.5<br>(100%) | 231.6 ± 143.4<br>(80%) | **6.45e-04** | LR < ES |
|  | Monkey W | 327.2 ± 96.3<br>(96%) | 188.5 ± 146.1<br>(74%) | **2.67e-09** | LR < ES |
|  | Monkey T | 234.1 ± 138.0<br>(90%) | 227.3 ± 148.3<br>(88%) | 7.44e-01 | LR ∼ ES |
| DLPFC | Pooled | 344.9 ± 27.5<br>(100%) | 272.4 ± 13.6<br>(100%) | **7.10e-34** | LR < ES |
|  | Monkey W | 259.4 ± 149.5<br>(59%) | 163.6 ± 141.6<br>(76%) | **3.89e-04** | LR < ES |
|  | Monkey U | 353.2 ± 81.3<br>(99%) | 363.1 ± 41.8<br>(100%) | 3.87e-01 | ES ∼ LR |
|  | Monkey T | 322.7 ± 100.8<br>(97%) | 274.0 ± 20.8<br>(100%) | **9.54e-17** | LR < ES |
| OFC | Pooled | 314.9 ± 20.1<br>(100%) | 269.6 ± 35.1<br>(100%) | **5.32e-21** | LR < ES |
|  | Monkey W | 364.3 ± 76.7<br>(96%) | 187.5 ± 151.2<br>(80%) | **1.70e-13** | LR < ES |
|  | Monkey U | 319.2 ± 23.7<br>(100%) | 263.9 ± 45.5<br>(98%) | **1.35e-24** | LR < ES |
|  | Monkey T | 248.2 ± 133.5<br>(79%) | 285.5 ± 116.1<br>(92%) | 7.02e-02 | ES ∼ LR |

**Supplementary Table 12.** Onset-latency comparisons between brain areas for good-based and action-based choice signals. Mean ± SD (ms), with valid percentage in parentheses, across 100 iterations for the earning/spending (ES) and left/right (LR) choice signals. Valid percentage is the percentage of the 100 iterations that yielded a latency estimate (see Methods, Latency). Two-sided Wilcoxon rank-sum tests compare brain areas; significant *p* values are bold. Latency ordering ranks the means; *<* denotes an earlier signal and ∼ a nonsignificant adjacent comparison, not equivalence. Ordering colors identify ACC, DLPFC and OFC in red, green and blue, respectively. ACC was not recorded in monkey U.

| Variable | Dataset | ACC | DLPFC | OFC | *p*(ACC, DLPFC) | *p*(ACC, OFC) | *p*(DLPFC, OFC) | Latency<br>ordering |
|---|---|---|---|---|---|---|---|---|
| ES | Pooled | 216.8 ± 96.8<br>(100%) | 215.9 ± 101.7<br>(100%) | 195.5 ± 63.2<br>(100%) | 4.37e-01 | **1.02e-03** | **1.38e-04** | OFC < DLPFC ∼ ACC |
|  | Monkey W | 240.7 ± 115.5<br>(93%) | 273.3 ± 119.3<br>(35%) | 246.3 ± 123.2<br>(92%) | 9.14e-02 | 6.08e-01 | 1.56e-01 | ACC ∼ OFC ∼ DLPFC |
|  | Monkey U | – | 242.6 ± 104.3<br>(96%) | 159.5 ± 84.6<br>(100%) | – | – | **1.67e-11** | OFC < DLPFC |
|  | Monkey T | 199.4 ± 122.7<br>(77%) | 212.9 ± 128.0<br>(94%) | 222.3 ± 121.2<br>(67%) | 4.29e-01 | 2.52e-01 | 8.75e-01 | ACC ∼ DLPFC ∼ OFC |
| LR | Pooled | 178.2 ± 135.6<br>(63%) | 143.0 ± 75.0<br>(100%) | 176.6 ± 72.3<br>(100%) | 6.99e-02 | 9.92e-01 | **2.67e-05** | DLPFC < OFC ∼ ACC |
|  | Monkey W | 168.9 ± 139.1<br>(41%) | 184.8 ± 148.1<br>(50%) | 195.1 ± 145.6<br>(69%) | 5.79e-01 | 3.13e-01 | 7.24e-01 | ACC ∼ DLPFC ∼ OFC |
|  | Monkey U | – | 223.8 ± 115.0<br>(100%) | 187.0 ± 64.1<br>(98%) | – | – | **9.89e-06** | OFC < DLPFC |
|  | Monkey T | 226.9 ± 131.1<br>(64%) | 143.6 ± 72.3<br>(100%) | 218.1 ± 112.1<br>(81%) | **7.99e-05** | 5.20e-01 | **5.14e-07** | DLPFC < OFC ∼ ACC |

**Supplementary Table 13.**
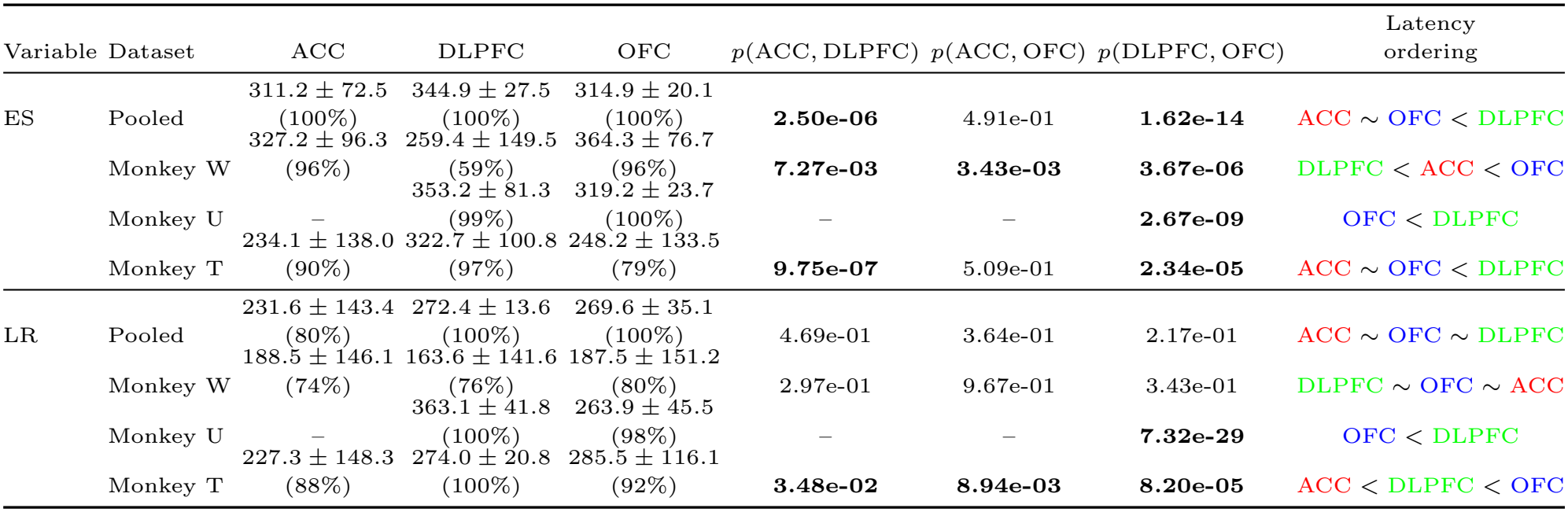
Half-peak latency (*t*_50_) comparisons between brain areas for good-based and action-based choice signals. ES and LR denote earning/spending and left/right choice signals, respectively. Same conventions as Supplementary Table 12, except that latency is quantified by *t*_50_.

